# Physiological and pathological changes in the enteric nervous system of rotenone-exposed mice as early biomarkers for Parkinson’s disease

**DOI:** 10.1101/2020.12.11.421164

**Authors:** Gabriela Schaffernicht, Qi Shang, Alicia Stievenard, Kai Bötzel, Yanina Dening, Romy Kempe, Magali Toussaint, Daniel Gündel, Mathias Kranz, Heinz Reichmann, Christel Vanbesien-Mailliot, Peter Brust, Marianne Dieterich, Richard H.W. Funk, Ursula Ravens, Francisco Pan-Montojo

## Abstract

Parkinson’s disease (PD) is known to involve the peripheral nervous system (PNS) and the enteric nervous system (ENS). Functional changes in PNS and ENS appear early in the course of the disease and are responsible for some of the non-motor symptoms observed in PD patients like constipation, that can precede the appearance of motor symptoms by years. We have shown that environmental toxins can trigger the disease by acting on the ENS and on the autonomic nervous system. Oral exposure to the pesticide rotenone, a mitochondrial Complex I inhibitor, leads to decreased stool depositions in mice. Here we analyzed the effect of rotenone on the function and structure of the ENS by measuring intestinal contractility in a tissue bath and by analyzing related protein expression. Our results show that rotenone changes the normal physiological response of the intestine to carbachol, dopamine and electric field stimulation. The magnitude and direction of these alterations varies between intestinal regions and exposure times and is associated with an early up-regulation of dopaminergic, cholinergic and adrenergic receptors and an irregular reduction in the amount of enteric neurons in rotenone-exposed mice. The early appearance of these alterations makes them ideal candidates to be used as biomarkers for the detection of Parkinson’s disease in its early stages.

## Introduction

Hallmark lesions of Parkinson’s disease (PD) were traditionally considered to be present in the dopaminergic neurons of the substantia nigra (SN) and in the noradrenergic neurons of the locus coeruleus (LC). However, pathological studies showed that PD patients typically have lesions in other central nervous system (CNS) and PNS structures (e.g. the ENS, the sympathetic coeliac ganglion (CG), the intermediolateral nucleus (IML) of the spinal cord, the motor nucleus of the vagus DMV or the amygdala) [1, 2]. These lesions are not exclusive of PD [3] and mainly consist in intraneuronal and intraglial alpha-synuclein aggregates called Lewy bodies (LB) and Lewy neurites (LN).

In idiopathic PD most patients showing PD-related inclusions in CNS sites also present LB and LN in the ENS and the sympathetic ganglia [4]. Based on autopsies performed on PD patients and healthy individuals, Braak and colleagues proposed a pathological staging of the disease [5]. According to this staging, PD lesions follow a spatio-temporal pattern that starts in the olfactory bulb (OB) and the ENS progressing into the CNS through synaptically connected structures. This pathological staging of the disease seems to correlate well with the appearance of early non-motor symptoms in PD patients including hyposmia, gastrointestinal alterations, autonomic dysfunction and pain [6].

Little is known about the degenerative process of the ENS in PD patients. Physiological studies in patients revealed delayed gastric emptying, dystonia of the external anal sphincter that causes difficult rectal evacuation and general slow transit constipation probably caused by the local loss of dopaminergic neurons [7–9]. In recent years many groups have studied expression and modification of enteric alpha-synuclein in PD patients, with controversial results. Whilst some found an increase in alpha-synuclein inclusions and phosphorylated alpha-synuclein in the above-mentioned areas when compared to control subjects [10], there appears to be a high variability between patients. A review of the literature reveals that not many studies investigated PD-associated changes in the function of the ENS, its neuronal composition or its extrinsic innervation.

In previous studies, we have shown that chronic oral administration of rotenone in mice induces alpha-synuclein accumulation in most of the regions described in Braak’s staging [11, 12]. The appearance of these alterations followed a spatio-temporal pattern similar to the one predicted by Braak and colleagues and the disruption of the autonomic nerves connecting the gut to the CNS (i.e. the vagus and sympathetic nerves) prevented the progression of the pathology. Interestingly, we also observed gastrointestinal alterations in rotenone-exposed mice in the form of constipation and a reduction of the noradrenergic innervation of the gut [13]. An independent recent study showed alterations in the extrinsic cholinergic innervation of the gut after rotenone exposure in guinea pigs [14]. In this context, the current study was designed to further investigate the consequences of orally administered rotenone on the ENS and on gastrointestinal contractility (GC). Our results show that chronic rotenone administration in mice affects the electrophysiological properties of the gut and that these alterations correlate well with the up-regulation of certain post-synaptic receptors, and that this is most probably due to the reduction of the extrinsic input from the sympathetic and vagal innervation. This could be important to identify biomarkers for the early diagnosis of PD.

## Materials and methods

### Animal model

#### Animal housing

1 year-old C57/BL6J mice (Janvier, France) were housed at room temperature under a 12-h light/dark cycle. Food and water were provided *ad libidum*. All animal experiments were carried out in accordance with the National Institutes of Health Guide for the Care and Use of Laboratory Animals, and protocols were approved by the Saxonian Committee for Animal Research in Germany.

### Chronic oral rotenone administration

Chronic oral exposure to low doses of the pesticide rotenone was performed as previously described (Pan-Montojo et al. 2010). Briefly, 1 year-old mice were divided into 2 groups (n= 10) and exposed 5 days a week for up to 4 months. A 1,2 mm x 60 mm gavage needle (Unimed, Switzerland) was used to administer 0.01 mL/g animal weight of rotenone solution corresponding to a 5 mg/kg dose. Controls were exposed only with the vehicle solution (2% carboxymethylcellulose (Sigma-Aldrich, Germany) and 1.25% chloroform (Carl Roth, Germany)).

### Tissue preparation for organ bath and Western blot

Mice were killed with an overdose of ketamine. Several 1 cm long adjacent intestinal segments were removed from each region (duodenum, jejunum, ileum and colon), submerged in Krebs solution (120 mM [NaCl], 5.8 mM [KCl], 2.5 mM [CaCl_2_], 1.2 mM [MgCl_2_], 1.4 mM [NaH_2_PO_4_], 15 mM [NaHCO_3_], 11 mM [glucose]), and gently flushed to remove luminal contents. The segments were used for contractility studies in an organ bath or immediately frozen in liquid nitrogen until protein extraction.

### Contractility studies in an organ bath

In order to test the motility of the intestinal segments we measured contractility using an isolated organ bath (Glasapparatemeister, Topas GmbH, Germany) as performed by others [15]. The tissue pieces were ligated at each end with a silk thread and suspended longitudinally between two platinum electrodes by attaching one end to the isometric force transducer (Transducer Typ SG4-45, SWEMA, Sweden) and the other end to the base of the 15 ml chamber filled with Krebs solution maintained at 37°C and aerated with 95% O_2_/5%CO_2_. Samples were stabilized in the Krebs solution for 45 minutes. During this time the bath solution was changed twice and samples were pre-loaded to 10 mN by step-wise increases until the tension was adjusted at 10 mN. Spontaneous smooth muscle activity (SMA) and responses to electrical field stimulation (EFS) were measured isometrically. The mechanical activity of the muscles was recorded using a transducer amplifier relayed to a bioelectric amplifier (Multiplexing Pulse Booster 3165, Gemonio-Varese, Italy) equipped with the data acquisition Chart 4.0TM (AD Instruments, Sydney, NSW, Australia). The preparations were allowed to stabilize for 45 min. During this period the Krebs solution in the bath was exchanged 2 times. After this equilibration period, SMA was characterized for 15 min by recording basal tone (minimum force during contraction), contraction amplitude (difference between maximum and minimum force during a contraction) and frequency of contractions (number of spontaneous contractions per minute). The preparations were then exposed to different concentrations (in Mol/L: 10^−6^, 3×10^−6^, 10^−5^, 3×10^−5^, 10^−4^, 3×10^−4^, 10^−3^) of the neurotransmitters dopamine and noradrenaline or to the unselective muscarinic receptor agonist carbachol, and the responses of the 3 contractile parameters were determined. After several medium changes, samples were subjected to EFS using an electric stimulator (Föhr Medical Instruments, Seeheim/Ober Beerbach, Germany). The optimal EFS parameters were determined testing several values for voltage (5, 10 and 15V) and frequencies (10, 16 and 80 Hz) on test samples. We decided to stimulate the ENS from 2-months exposed mice using three different pulse durations (200, 400 and 800 μs) and reduced the number to two different pulse durations (200 and 400 μs) for the 4-months exposed mice. In all cases a train duration of 30 s with a voltage of 10V, a pulse delay of 0 s and a pulse rate of 16Hz were used. Responses to EFS were determined after the last change of bath solution in absence of any additional drugs and represented the responses to stimulation of ENS alone. In order to study the contribution of various neurotransmitters in ENS activation the following drugs were added in top of each other: atropine (ATR, 10^−5^Mol/L) for blocking cholinergic receptors, plus guanethidine (GUA, 10^−5^Mol/L) to block additional noradrenergic receptors, plus serotonin (SHT, 10^−4^Mol/L) to analyse the effect of serotonin in the absence of cholinergic and adrenergic inputs and plus tetrodotoxin (TTX, 10^−6^Mol/L) to additionally block voltage-gated sodium channels.

Contractility data for each time-point was translated into a curve by the data acquisition system. The contractile parameters maximum force, minimum force, amplitude and frequency of spontaneous contractions before and after each exposure were automatically extracted using a self-made program written with Matlab (version R2014b, Mathworks, Massachusetts, USA). The effects for each drug concentration on the contractile parameters were determined by normalizing the drug-induced changes to pre-drug control values (time 0). These values were used to generate concentration-response curves for each parameter.

### Protein extraction for Western blot

Protein extraction was done in an ice-cold Eppendorf holder using ice cold protein extraction buffer (60 mMol/L TrisHCl (pH 6.8), 1mMol/L Na_3_VO_4_, 1% SDS) (300μL). Intestine samples (n=5 per experimental group) were cut into smaller pieces, homogenized with the help of a homogenizing pestle in 1,5 mL Eppendorf tubes and an ultrasound tip. The homogenized sample was kept at 99°C for 5 minutes and centrifuged at 1400 rpm for 5 minutes. The supernatant was collected and stored at −80 °C until use.

### SDS-PAGE and Western blotting

Protein concentration from intestine extracts was measured using the Nanodrop ×100. Samples were then diluted in each case to have the same end protein concentration of 30μg in 20 μL and protein extract, and were loaded together with 5 μL of loading buffer (Invitrogen, Germany, EU) into a 12% polyacrylamide gel.

Page ruler 10-250 kDa molecular weight marker was used for size estimation. Using a Mini Gel Tank (Life Technologies, USA), proteins were separated by electrophoresis using 80V (stacking) and 120V (migration) and transferred onto a nitrocellulose membrane using 10V for 3 h. To verify the efficiency of the transfer, membranes were incubated in a Red Ponceau solution during 2 min and washed 3 times (3 x 5min) in deionized water. Membranes were blocked in a blocking solution containing Tris-buffered saline + 0.1% Tween 20 (TBST 1X) and 5% (w/v) non-fat dry milk for 1 hour. Membranes were then incubated overnight at 4°C with one of the following primary antibodies: rabbit anti-tyrosine hydroxylase (1:1000, Pel Freez), mouse anti-GAPDH (1:1000, Thermo Scientific), rabbit anti-ChAT (1:1000, Abcam), rabbit anti-Muscarinic Acetylcholine Receptor (M3, 1:3000, ab126168, Abcam), rabbit anti-Dopamine receptor D2 antibody (ADI-905-740-100, 1:1000, Enzo Life Science) or rabbit anti PGP9.5 (AB1761, 1:1000, Millipore or 1:3000, Enzo life Science). On the following day, blots were washed 3 x 5 min. with TBST 1X and incubated with horseradish peroxidase-conjugated anti-rabbit or anti-mouse secondary antibodies (1:5000, Abcam) at room temperature for 1 h.

The blots were subsequently washed 3 x 5 min in TBST 1X and developed with the Prime Western Blotting detection Reagent (GE Healthcare Amersham) using the LAS3000 Bioimager (Fujifilm, Germany, EU). Images were then processed using the open access ImageJ based program FIJI (ww.fiji.sc), where only minor adjustment of the brightness and contrast were performed.

### Small animal PET/MR imaging

PET studies of the gastrointestinal tract were performed in male C57/BL6J mice in two experimental series. The first experimental series (age = 14 months; weight = 28–35 g), was composed of a control group (n = 5) and a treated group (n = 6) respectively unexposed or exposed for 2 months to rotenone. The second experimental series (age = 16 months; weight = 28–35 g), was composed of a control group (n = 2) and a treated group (n = 5) respectively unexposed or exposed for 4 months to rotenone. For dynamic PET studies the animals received an intravenous injection of (−)-[^18^F]Flubatine into the tail vein (6.4 ± 1.9 MBq; 0.63±0.58 nmol/kg ; A_m_ : 1016±712 GBq/μmol (EOS) for the 2 months treated serie) or retro-orbital (6.5 ± 2.4 MBq; 0.18±0.18 nmol/kg; A_m_ : 4460±2115 GBq/μmol (EOS); for the 4 months treated serie) followed by a 60-min PET/MR scan (Mediso nanoScan®, Budapest, Hungary). A T1-weighted WB gradient echo sequence (GRE, repetition time = 20 ms, echo time = 6.4 ms) was performed for AC and anatomical orientation. The reconstruction parameters for the list mode data were the following: 3D-ordered subset expectation maximization (OSEM), 4 iterations, 6 subsets. The mice were positioned prone in a special mouse bed (heated up to 37°C), with the head fixed to a mouth piece for the anesthetic gas supply with isoflurane in 40% air and 60% oxygen (anesthesia unit: U-410, Agnthos, Lidingö, Sweden; gas blender: MCQ, Rome, Italy), image co-registration and evaluation of the volume of interest (VOI) were performed with PMOD (PMOD Technologies LLC, v. 3.9, Zurich, Switzerland). Spherical VOI with diameters of 1 to 2 mm were placed at the center of the caecum on the PET/MR fusion image. For the jejunum the VOI was delineated from the PET signal. The activity concentrations in the VOIs are expressed as the mean standardized uptake values (SUV) +/− SEM.

### Statistical analysis

Concentration-response curves were compared using the 2-way ANOVA test with a Bonferroni’s post-hoc test. Contractility data regarding frequency and EFS were analyzed using an unpaired Student t-Test. Protein expression data was analyzed using an unpaired non-parametric Student t-test (Mann-Whitney Test). Significance was set at p<0.05. The time-activity curves were analysed using an unpaired Student t-Test. Significance was set at p<0.05.

## Results

### Rotenone exposure induces changes in gastrointestinal contractility in response to carbachol and dopamine

The effect of rotenone on the cholinergic and dopaminergic inputs to the ENS was determined by comparing the effects carbachol and dopamine on motility of intestinal tissue from vehicle and rotenone-exposed mice. Our results show that rotenone induces region- and exposure time-dependent changes in the concentration-response curves for carbachol and dopamine. We did not observe any effects on the basal frequency or mean basal contraction amplitude (i.e. after equilibration the samples and before adding any drugs) between intestinal segments from vehicle-(control) and rotenone-exposed mice (Supplementary Figure 1).

#### Changes in the response to carbachol after 2 and 4 months of rotenone exposure

Carbachol is a cholinergic receptor agonist that enhances intestinal contractility. We constructed concentration-response curves for carbachol in all intestinal segments from control and rotenone-exposed mice (2 months and 4 months of exposure) (see Figure 1).

**Figure 1:**
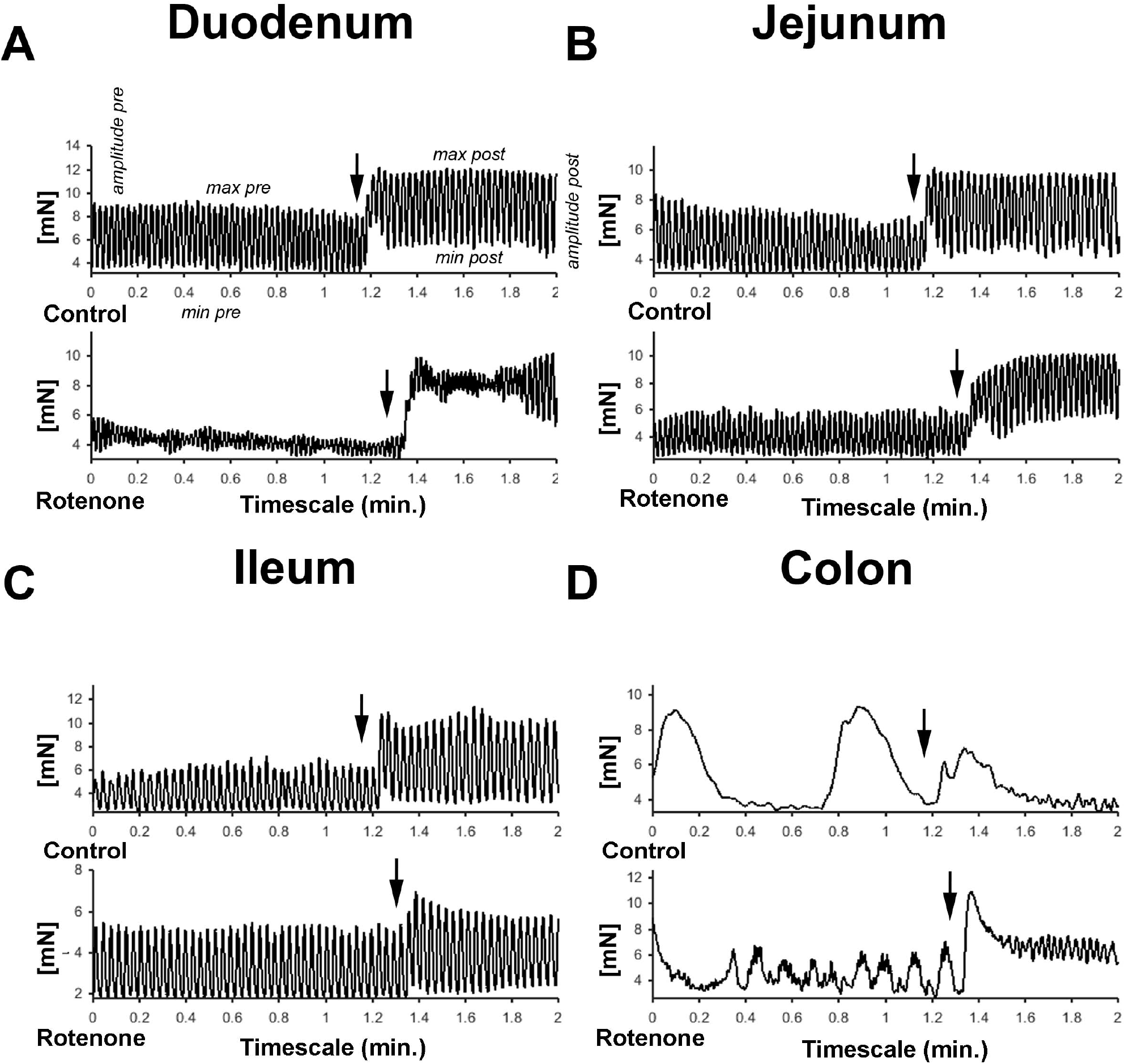
**A-D** graphic showing representative examples of the force generated by the peristaltic movements and the effect of carbachol in the different gut segments in mN. Arrows in all graphics indicates the moments carbachol was added to the buffer.

In comparison with intestinal tissue from unexposed mice, 2 months of rotenone exposure led to larger increases in the maximum force in response to carbachol in duodenum and jejunum (see Supplementary Figure 2A and E), to elevation of carbachol-stimulated tone in duodenum (see Supplementary Figure 2B) and higher contraction amplitudes in the duodenum, jejunum and ileum (see figure 2A, B and C). In comparison to vehicle exposure, 4 months of rotenone exposure produced a slightly different pattern of changes: carbachol increased maximum contractility to a larger extent in the jejunum and to a lower extent in the ileum (see Supplementary Figure 2G and K), increased tone in the jejunum (see Supplementary Figure 2H) to a larger extent, and induced larger contraction amplitudes in the jejunum and the colon (see Figure 2F and H and table 1)

**Figure 2:**
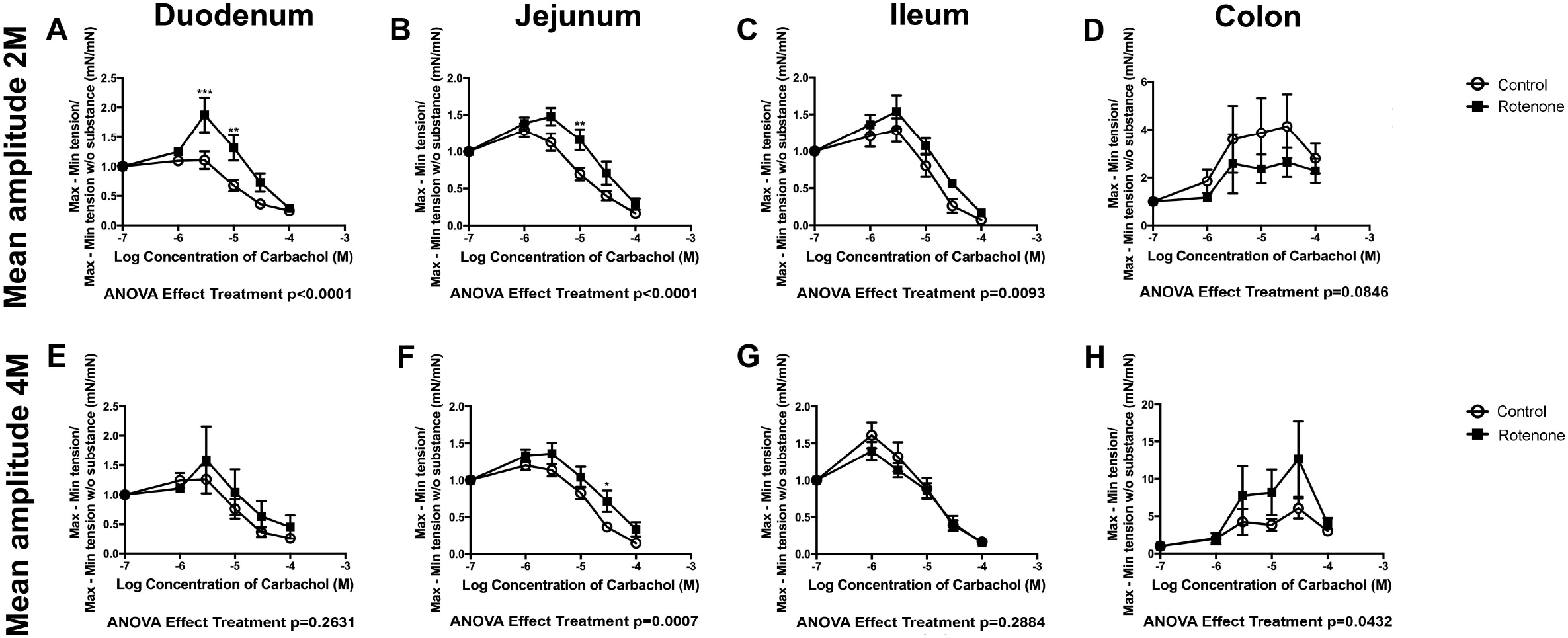
2- and 4-Month rotenone exposure in mice increases the sensibility to carbachol in duodenum and jejunum. Effect of carbachol treatment on the amplitude (Mean amplitude) of the duodenum (**A and E**), jejunum (**B and F**), ileum (**C and G**) and colon (**D and H**) from control or 2 or 4 months rotenone-exposed mice. Data was analyzed using a 2-way ANOVA and a Bonferroni post-hoc test. Values of the effect of exposure are shown under the corresponding graph. *, **, *** correspond to *P*<0.05, *P*<0.01 and *P*<0.001 respectively for each concentration obtained with the Bonferroni post-hoc test. Error bars represent SEM in all graphics.

#### Changes in response to dopamine after 2 and 4 months of rotenone exposure

We then analyzed the effect of rotenone on the responses to dopamine among the different regions of the intestine. The neurotransmitter dopamine induces intestinal relaxation by decreasing tone, maximum force development and amplitude of contraction. (see Figure 3) Two months of rotenone exposure led to a bigger dopamine-induced decrease in tone of duodenum, jejunum and ileum than in control preparations (see Supplementary Figure 3B,F and J). We did not observe any significant difference in the effect of dopamine on the maximum force of contraction or on contraction amplitude after 2 months of exposure with rotenone in comparison with vehicle exposure. However, after 4 months of exposure to rotenone, dopamine induced bigger decreases in the maximum force of contraction in jejunum and ileum (see Supplementary Figure 3G and K), larger relaxation of tone in duodenum and ileum (see Supplementary Figure 3D and L) and greater depression of mean contraction amplitude in duodenum, jejunum and ileum (see Figure 4 E, F and G). Interestingly, the effects in colon were the opposite direction with smaller increase in the maximum force of contraction and a weaker relaxation in tone of rotenone-exposed samples (see Supplementary Figure 3O and P and Table 2 for all results)

**Figure 3:**
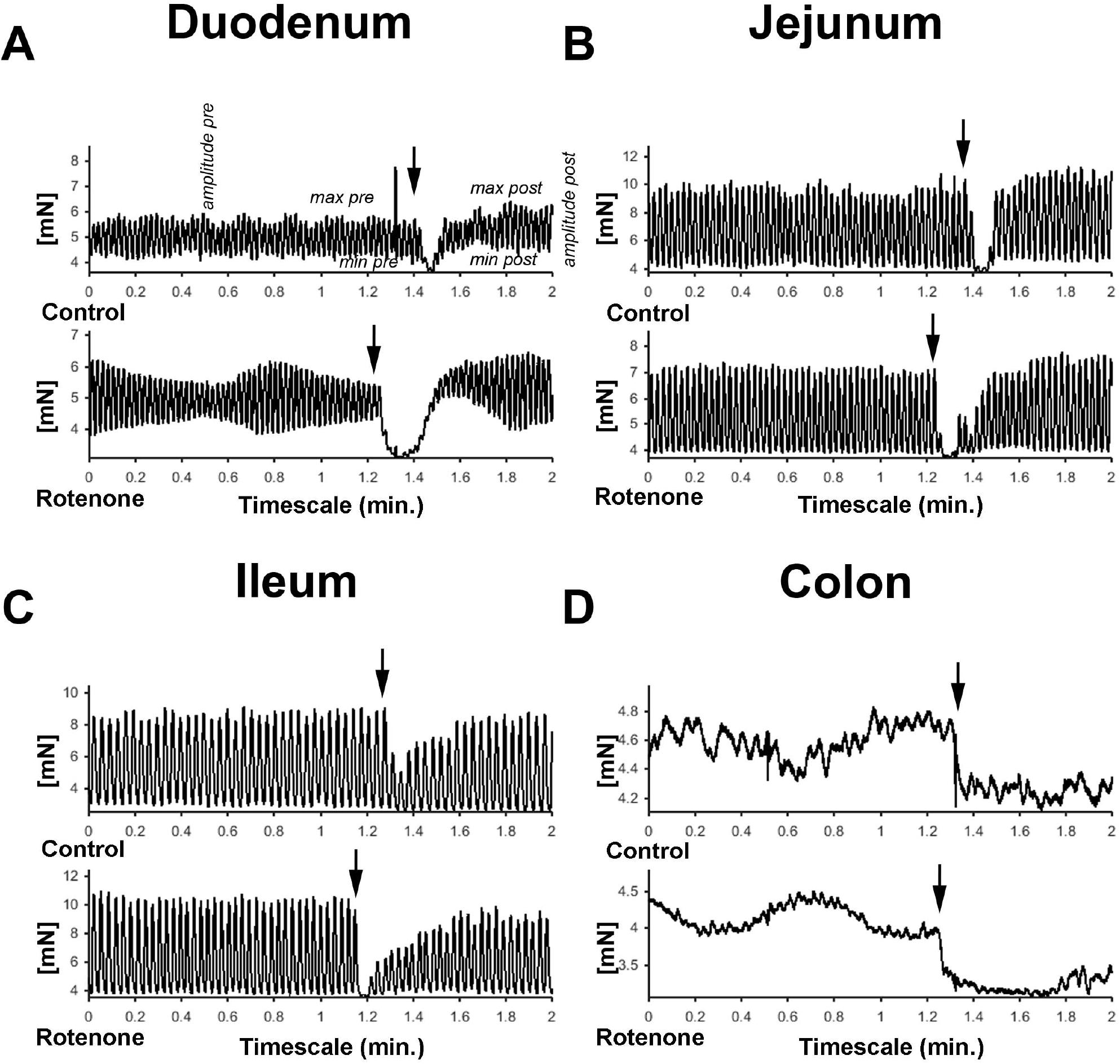
**A-D** graphic showing representative examples of the force generated by the peristaltic movements and the effect of dopamine in the different gut segments in mN.

**Figure 4:**
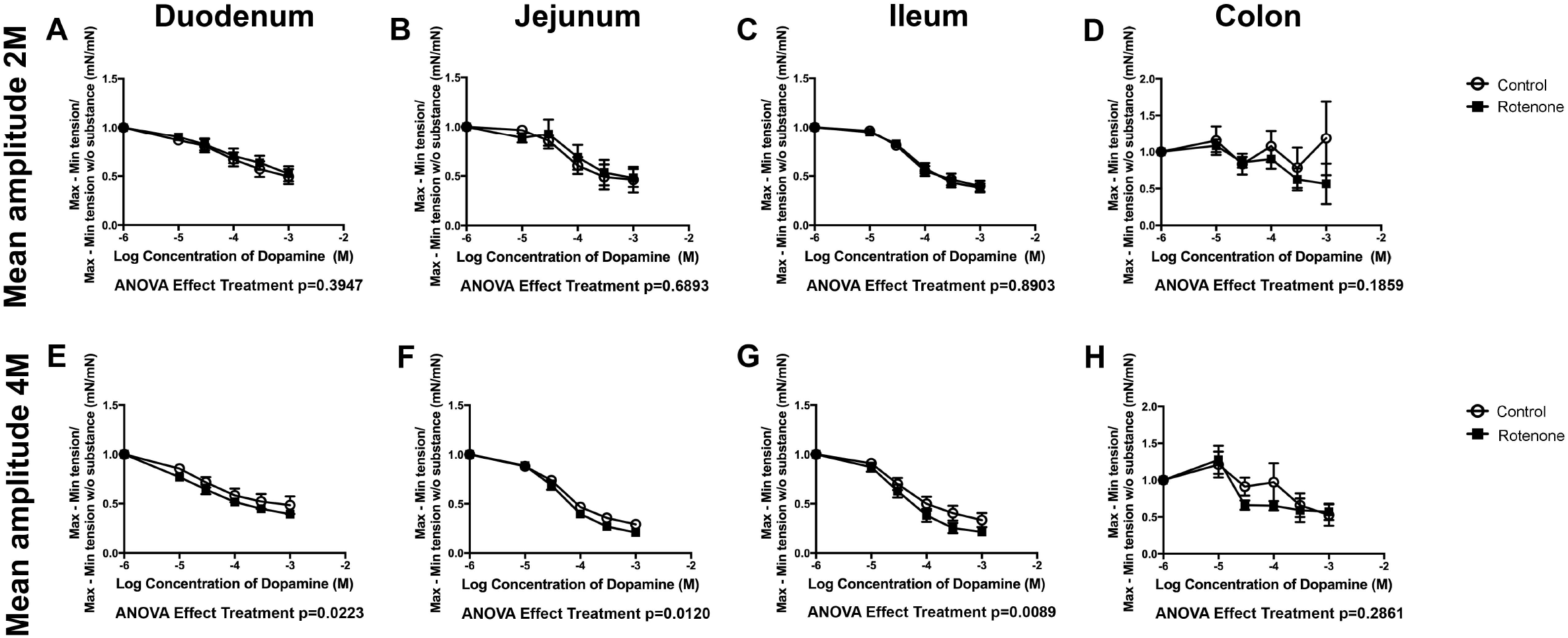
2- and 4-Month rotenone exposure in mice increases the sensibility to dopamine in duodenum and jejunum. Effect of dopamine treatment on the amplitude (Mean amplitude) of the duodenum (**A and E**), jejunum (**B and F**), ileum (**C and G**) and colon (**D and H**) from control or 2 or 4 months rotenone-exposed mice. Data was analyzed using a 2-way ANOVA and a Bonferroni post-hoc test. Values of the effect of exposure are shown under the corresponding graph. *, **, *** correspond to *P*<0.05, *P*<0.01 and *P*<0.001 respectively for each concentration obtained with the Bonferroni post-hoc test. Error bars represent SEM in all graphics.

Taken together, these results show that in general in mice exposed to rotenone carbachol and dopamine have increased effects when compared to the control group, suggesting an over-expression of the postsynaptic receptors in ENS neurons due to a decreased parasympathetic and dopaminergic input. From a physiological point of view these effects are expected to be quite variable and even divergent between intestinal regions and the duration of the exposure. However the signal to noise background ratio is still above the threshold. Such a reduction in the cholinergic input would be in accordance with previous observation from our lab and could be responsible for decreased intestinal motility previously observed using the 1-hour stool collection test [13].

### Rotenone exposure induces changes in the response of the ENS to electric field stimulation

We then tested the effect of rotenone exposure on the response of the ENS to electric field stimulation (EFS). EFS allows to induce indirect contractile effects on the intestine due to excitation of ENS and subsequent neurotransmitter release without directly electrically stimulating the intestinal smooth muscle. The response to EFS depends on the distribution of the neuronal subtypes within the different various regions of the intestine (i.e. cholinergic, serotoninergic, dopaminergic, nitrergic, etc. neurons). Using the parameters chosen for the experiments (see Material and Methods) EFS with a pulse duration of 200 (EFS 200) and 400 (EFS 400) μs (and using the parameters described in Material and Methods) produced a biphasic TTX-sensitive response in control tissue from non-exposed mice as reported previously [16]. This response consisted in an initial relaxation (R1) after the start of EFS followed by a contraction (C-Off) when the EFS stopped (see Figure 5).

**Figure 5:**
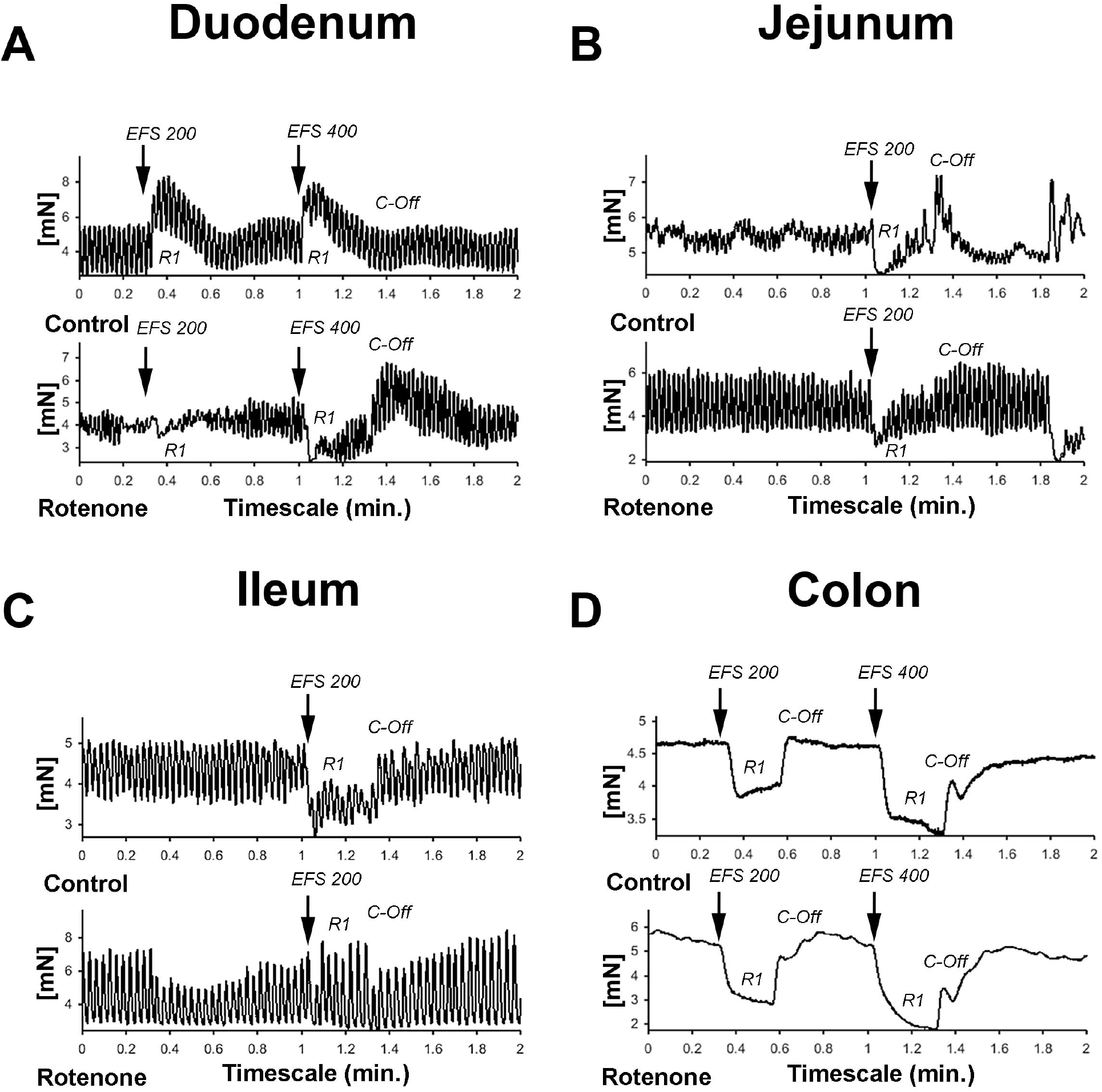
**A-D** graphic showing representative examples of the force generated by the peristaltic movements and the effect of EFS in the different gut segments in mN. Electric field stimulation normally induces an initial relaxation (R1) followed by a contraction (C-Off) when the stimulus ended.

#### Four months but not 2 months rotenone exposure increases the response of jejunum and ileum to EFS in the absence and presence of atropine, guanethidine or serotonin

In order to compare the responses to EFS in control and rotenone-exposed mice, we performed EFS 200 and EFS 400 in the absence or in the presence of atropine (AT), AT and guanethidine (G) (2 months) or AT, G and serotonin (S) (4 months). We then compared both curves using a 2-way ANOVA. We could not detect any significant differences during EFS 200 or EFS 400 stimulation between control and 2 months rotenone-exposure (see Supplementary Figure 4A-H and 5A-H). After 4 months of rotenone exposure intestinal muscle relaxations were enhanced during EFS 200 stimulation (R1 phase) in the ileum (see Supplementary Figure 4M) and so were reactive contractions when stimulation stopped (C-Off phase) in the jejunum and ileum (see Supplementary Figure 4L and N). Similar differences between exposures were observed in these regions when an EFS 400 was applied (see Supplementary Figure 5 L-N and Tables 3 and 4 for a summary)

Altogether, these results show that rotenone exposure leads to a stronger net inhibitory outcome of the enteric neuronal network thus suggesting either a stronger inhibitory or a weaker excitatory neuronal input in the ENS of rotenone-exposed mice. This data correlate well with the previously reported reduction in pellet output after 1,5 months of such a regimen (ie. chronic low dose rotenone exposure)[13]. In order to differentiate between these two options, we compared the effect of EFS alone and in the presence of i) atropine (AT), a competitive muscarinic acetylcholine (Ach) receptor antagonist to investigate non-cholinergic responses and ii) AT and guanethidine (G), a selective inhibitor of noradrenaline release from postganglionic adrenergic neurons to investigate non-adrenergic non-cholinergic responses. Additionally, we decided to analyze iii) AT, G and serotonin (SHT), a neurotransmitter that increases gut motility and the peristaltic reflex by inhibiting dopaminergic neurons among other effects, to investigate the basal levels of serotonin in the ENS.

### Rotenone exposure reduces the amount of acetylcholine released in duodenum, ileum and colon and increases it in the jejunum during EFS

To indirectly determine the amount of ACh released during the EFS, we compared the relaxation and contraction during the R1 and C-Off phases respectively in the absence and the presence of 10^−5^Mol/L of AT.

In the duodenum, we did not observe differences in the effect of AT on the R1-relaxation between controls and exposed mice after 2 months of rotenone exposure (see Figures 6A and 7A). The inhibitory effect of AT on the C-Off contraction induced by EFS 200 and EFS 400 was slightly stronger on exposed mice when compared to control mice (see Figures 6B and 7B). 4 months of rotenone exposure did not change the effect of AT on the R1-relaxation between controls and exposed mice (see Figures 6I and 7I), but the effect of AT on the C-Off contraction induced by EFS 200 and EFS 400 on exposed mice was stronger than on control mice (see Figures 6J and 7J and Tables 3 and 4 for a data summary).

**Figure 6:**
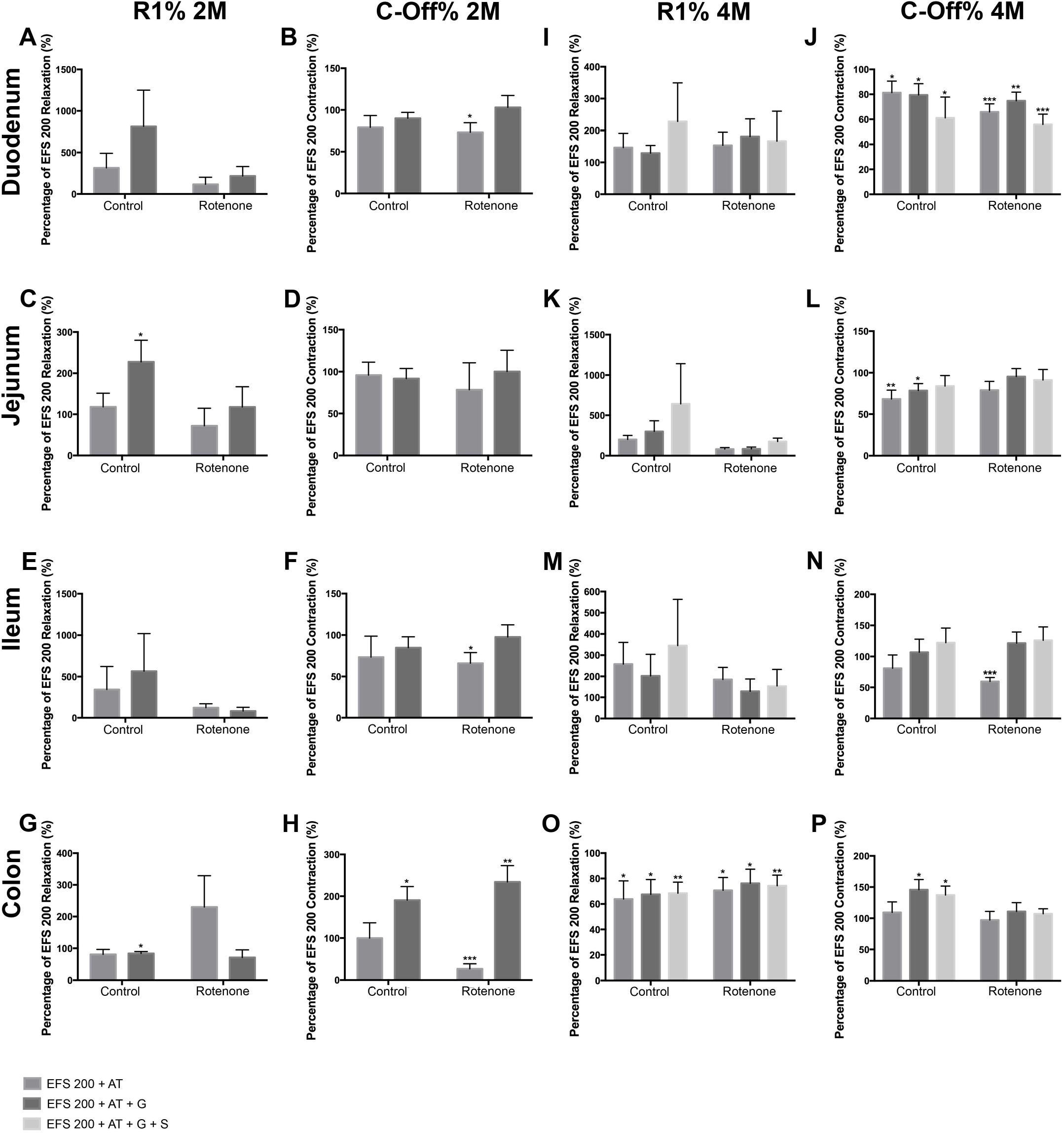
Effect of a 200 μs electric field stimulation (EFS 200) on the different intestinal regions after 2 and 4 months of rotenone exposure in mice. In all regions, AT induced a deeper relaxation and a lower contraction when compared to the reaction observed without atropine (AT) or guanethidine (G) when EFS 200 and EFS 400 were applied. Data in figures is shown as a percentage of the EFS 200 or EFS 400 relaxation/contraction obtained without neurotransmitters and the p values refer to this percentage, if an additional significant difference between control and exposed samples is shown, the p value will be written separately. (**A-H**) Graphics showing the R1 and the C-Off induced by EFS 200 on the duodenum (**A** and **B**), jejunum (**C** and **D**), ileum (**E** and **F**) and colon (**G** and **H**) in Krebs solution alone, in the presence of atropine or in the presence of atropine and guanethidine after 2 months of rotenone exposure. (**I-P**) Graphics showing the R1 and the C-Off induced by EFS 200 on the duodenum (**J** and **J**), jejunum (**K** and **L**), ileum (**M** and **N**) and colon (**O** and **P**) in Krebs solution alone, in the presence of atropine or in the presence of atropine and guanethidine after 4 months of rotenone exposure. Data was analyzed using a 2-way ANOVA Values of the effect of exposure are shown under the corresponding graph. *, **, *** correspond to *P*<0.05, *P*<0.01 and *P*<0.001 respectively. Error bars represent SEM in all graphics. Data was analyzed using a Student’s t-test, *, **, *** correspond to *P*<0.05, *P*<0.01 and *P*<0.001 respectively. Error bars represent SEM in all graphics.

**Figure 7:**
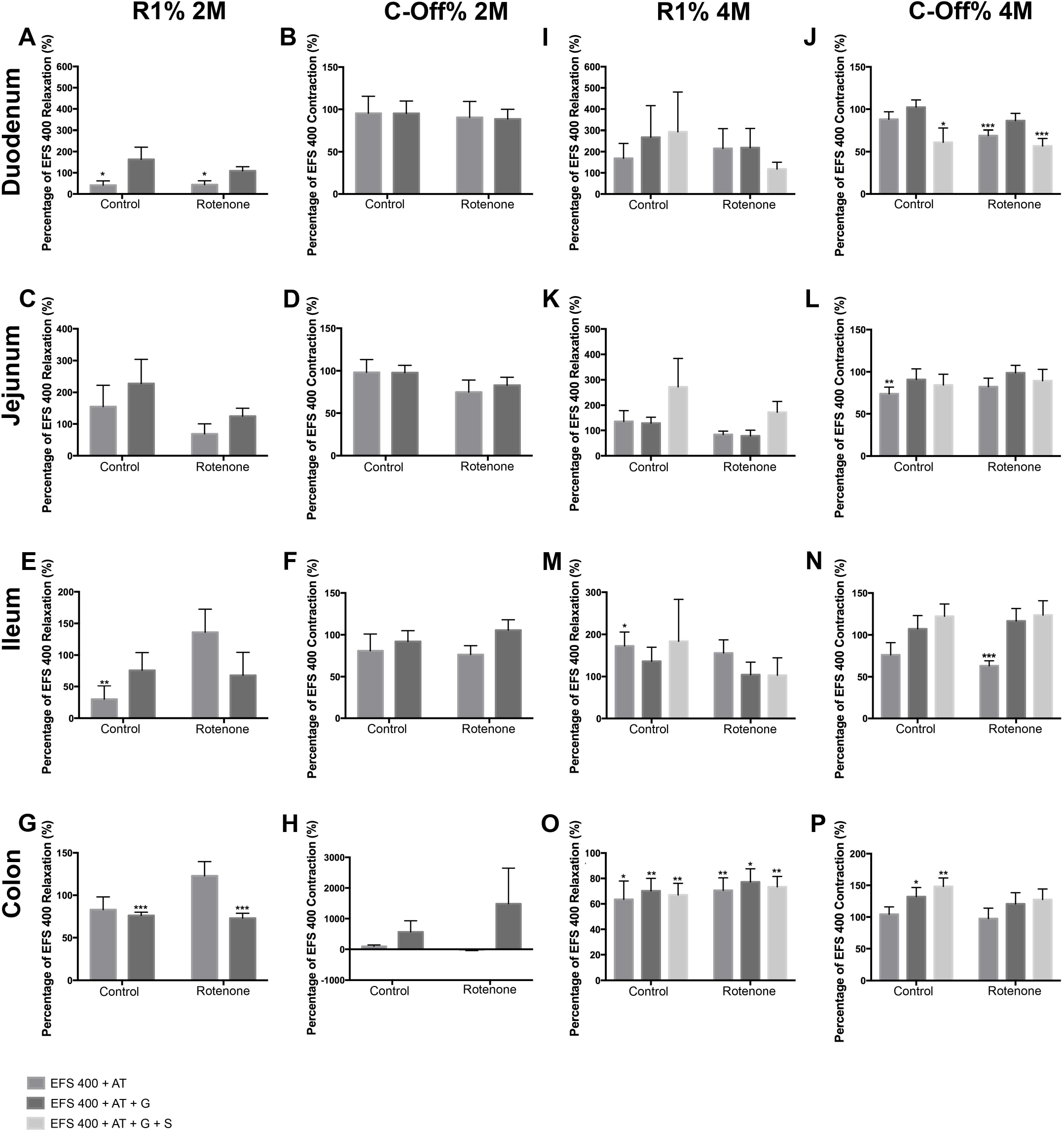
Effect of a 400 μs electric field stimulation (EFS 400) on the different intestinal regions after 2 and 4 months of exposure. In all regions, AT induced a deeper relaxation and a lower contraction when compared to the reaction observed without atropine (AT) or guanethidine (G) when EFS 200 and EFS 400 were applied. Data in figures is shown as a percentage of the EFS 200 or EFS 400 relaxation/contraction obtained without substances and the p values refer to this percentage, if an additional significant difference between control and exposed samples is shown, the p value will be written separately. (**A-H**) Graphics showing the R1 and the C-Off induced by EFS 200 on the duodenum (**A** and **B**), jejunum (**C** and **D**), ileum (**E** and **F**) and colon (**G** and **H**) in Krebs solution alone, in the presence of atropine or in the presence of atropine and guanethidine after 2 months of rotenone exposure. (**I-P**) Graphics showing the R1 and the C-Off induced by EFS 200 on the duodenum (**J** and **J**), jejunum (**K** and **L**), ileum (**M** and **N**) and colon (**O** and **P**) in Krebs solution alone, in the presence of atropine or in the presence of atropine and guanethidine after 4 months of rotenone exposure. Data was analyzed using a 2-way ANOVA Values of the effect of exposure are shown under the corresponding graph. *, **, *** correspond to *P*<0.05, *P*<0.01 and *P*<0.001 respectively. Error bars represent SEM in all graphics. Data was analyzed using a Student’s t-test, *, **, *** correspond to *P*<0.05, *P*<0.01 and *P*<0.001 respectively. Error bars represent SEM in all graphics.

In the jejunum, we observed no differences in the effect of AT on the R1 or C-Off phases of EFS 200 or EFS 400 after 2 months of exposure (see Figures 6C-D and 7C-D). 4 months after exposure, the effect of AT on R1-relaxation induced by EFS 200 and EFS 400 was significantly different between control and exposed samples (p<0.05). AT induced larger R1-relaxation on control samples and reduced the R1-relaxation in exposed samples (see Figures 6K and 7K). The differences observed on the effect of AT on the C-Off phase between rotenone exposed and control samples was opposite to the one observed in the duodenum, the ileum and the colon. AT induced a significant decrease in the contractility of control mice, whereas the decrease observed on exposed mice was smaller and non-significant (see Figures 6L and 7L and Tables 3 and 4).

In the ileum, 2 months exposure induced a higher response to AT (p<0.05) in the R1 phase with higher relaxations during EFS 400 in rotenone-exposed mice when compared to control samples (see Figure 7E). We also observed a significant reduction of the C-Off contraction in the presence of AT with the EFS 200 on exposed samples but not in controls (see Figure 7F). 4 months of rotenone exposure induced a slightly higher R1-relaxation in the presence of AT on control mice than in exposed mice (see Figures 6M and 7M). The effect of AT on the C-Off contractility was higher on exposed samples than controls (see Figures 6N and 7N and Tables 3 and 4).

In the colon EFS 200 and EFS 400 induced an opposite reaction to the one observed in all other intestinal segments in rotenone-exposed samples with a contraction in the R1 phase instead of a relaxation (see Figures 6G and O and 7G and O). The contraction during the R1 reaction was opposite and significantly different (p<0.05) to the relaxation observed in control samples during the EFS 200 (see Figure 6G). Additionally, we also observed an increased effect of AT on the C-Off phase during the EFS 200 with a significant reduction of the C-Off contraction in the presence of AT on exposed samples but not in controls (see Figure 6H). After 4 months of exposure, AT had no effect on the C-Off contractility in control or exposed mice (see Figures 6P and 7P). Interestingly, the effect of AT on the R1 EFS-induced relaxation after 4 months was opposite to all other intestinal regions and to the effect after 2 months of exposure. AT diminished the relaxation induced by EFS in both control and exposed samples (see Figures 6O and 7O). No difference between exposures was observed. (See Tables 3 and 4).

Overall the increased relaxation in the R1 phase and reduced percentage of contraction in the C-off phase suggest that also the amount of ChAT intrinsic to the enteric nervous system is also reduced in rotenone exposed mice when compared to control mice.

### 2 months but not 4 months rotenone exposure influences the noradrenergic, noncholinergic (NANC) reaction to EFS 200 or EFS 400

We then analyzed the NANC reaction of the different regions to the EFS 200 and EFS 400. For this we added 10^−5^M guanethidine to the Krebs solution, which already contained AT. As expected when inhibiting the noradrenergic input to the ENS, the NANC reaction to EFS showed non-significant increases in the R1 relaxation and non-significant higher C-Off contractions in most of the regions when compared to the reaction with AT at any time point in control mice. After 2 months of rotenone exposure, the only region showing a significant difference between the reaction to AT alone and to AT+G was the colon. In exposed samples but not in controls, the addition of guanethidine induced an increased contraction in the C-Off phase during EFS 200 and a reduced relaxation in the R1 phase during EFS (see Figure 7 G and H). After 4 months of exposure, we only observed a significant increase in the C-Off contraction after addition of guanethidine during EFS 200 and EFS 400 in the ileum of rotenone-exposed mice, the same region where the effect of AT had been the highest (see Figures 6P and 7P). Our results showed no significant differences in the NANC reaction to EFS 200 and EFS 400 between rotenone exposed and control samples in all other regions studied. (See Tables 5 and 6).

### ENS Reaction to Serotonin is not altered by rotenone exposure

In PD, the serotoninergic system in the brainstem is affected during the course of the disease. Therefore, we tested whether rotenone exposure affected the serotoninergic system in the gut. For this we added 10^−4^M of SHT in the presence of AT and G. In all regions and exposure groups except the colon, SHT addition induced an increased contractility with higher contractility forces. The magnitude of this increase was region dependent and no difference was observed between control and rotenone-exposed mice. In both groups the maximal contractility force was significantly higher (p<0.05) in the duodenum when compared to the ileum and the colon and the maximal contractility force in the jejunum was significantly higher to the one exerted in the colon (p<0.05) (see Figure 8A-F). The effect of SHT on the intestinal tonus depended on the region. It induced an increase of tonus in the duodenum and the jejunum, it had no effect on the colon and it decreased the tonus in the ileum. We did no observe any differences between rotenone-exposed and control samples (p>0.05) (see Figure 8A-F). (See Table 7 and 8)

**Figure 8:**
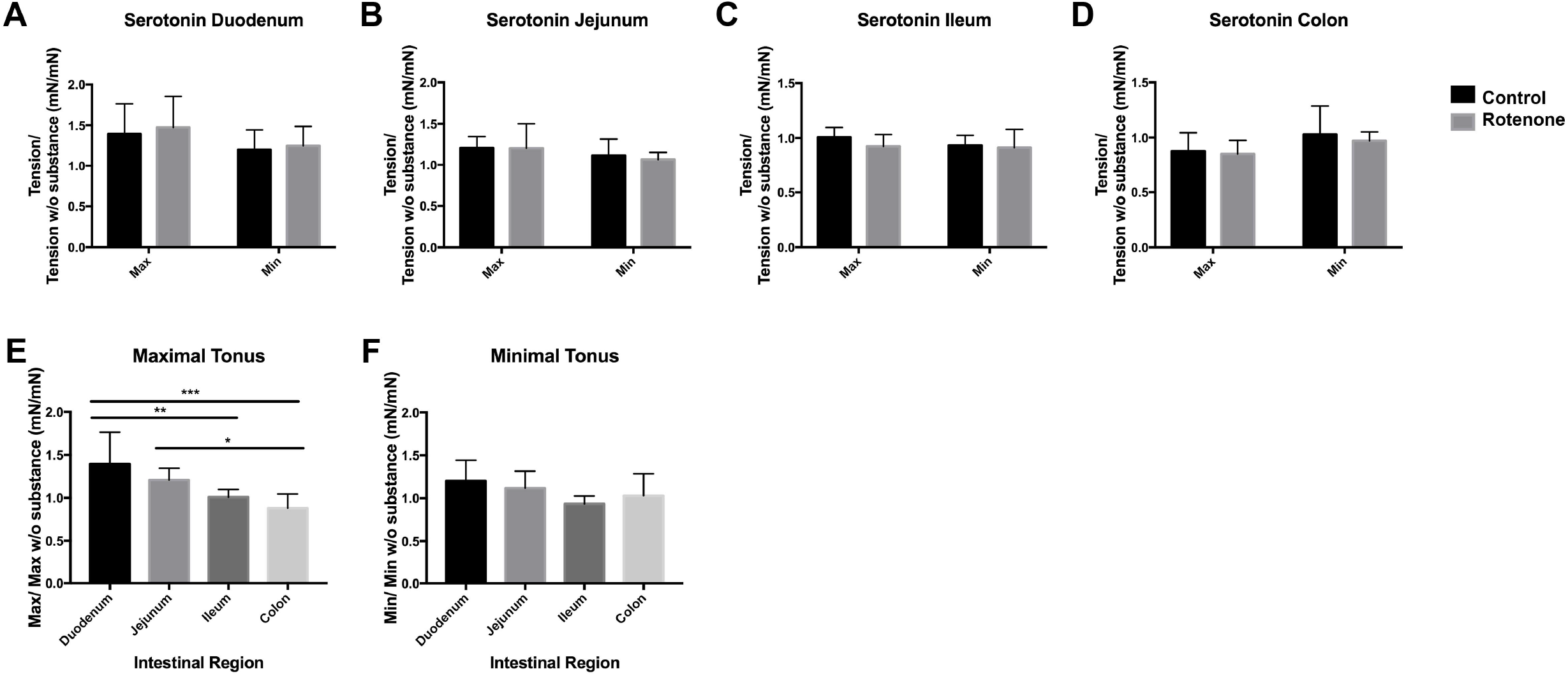
Effect of serotonin (SHT) on the maximal contractility force and the tone on all intestinal regions. (**A-D**) Bar graphs showing the effect of SHT on the duodenum (**A**), jejunum (**B**), ileum (**C**) and colon (**D**) of control (black) and rotenone-exposed (grey) mice. Bar graph in **E** shows a comparison of the maximal contractility force exerted by SHT between intestinal regions in control mice. Bar graph in **F** shows a comparison of the effect of SHT on the tone between intestinal regions in control mice. Data was analyzed using a Student’s t-test. , *, **, *** correspond to *P*<0.05, *P*<0.01 and *P*<0.001 respectively. Error bars represent SEM in all graphics.

Altogether, our results on contractility show that, in general, intestine sections from rotenone-exposed mice have an increased sensibility to carbachol and dopamine. Such increases are normally due to an up-regulation in the expression of the corresponding receptors as a result of a reduced neurotransmitter release, either due to alterations in the synapse or to the degeneration of the producing neurons. EFS experiments also suggest that the amount of acetylcholine might be reduced in the ENS, which could be caused by a reduction in the number of cholinergic neurons. In two previous studies, it has been shown that the cholinergic and adrenergic innervation of the gut is significantly reduced after rotenone exposure in mice and guinea pig [13, 14]. Therefore, we investigated the effect or rotenone exposure both on the expression of different postsynaptic receptors and on the enteric neuronal subpopulations.

### Cholinergic, adrenergic and dopaminergic postsynaptic receptors are up regulated in rotenone-exposed mice when compared to controls

First we compared the expression of muscarinic cholinergic- (M3), dopaminergic- (D2R) and adrenergic- (ß2-subunit) receptors between rotenone-exposed and control mice normalized to the total amount of neurons (PGP9.5) (n=5). These values were then normalized to the mean value in the control group. We could observe a region- and time-dependent up-regulation of all three kinds of receptors in rotenone-exposed mice. The expression of M3-muscarinic cholinergic receptors was up regulated in duodenum, jejunum, ileum and colon 2 and 4 months after rotenone exposure. The expression of adrenergic-(ß2-subunit) receptors was also up-regulated in duodenum, jejunum, ileum and colon after rotenone exposure. Likewise, the expression of D2-dopaminergic-receptors was generally up-regulated in the duodenum, jejunum, ileum and colon in rotenone-exposed mice (see Figures 9 and 10 and Tables 9, 10 and 11)

**Figure 9:**
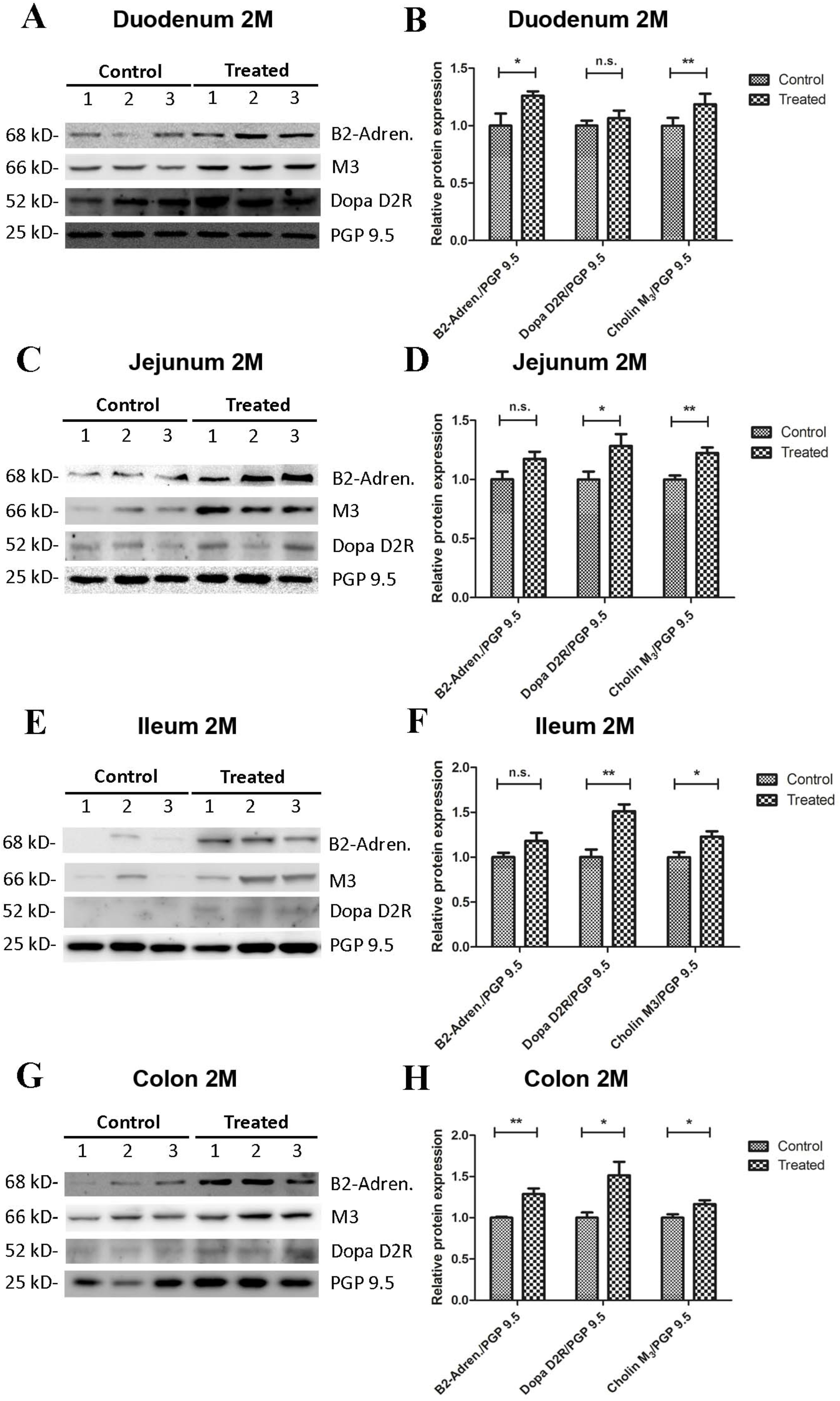
2-Month rotenone exposure induces an up regulation of the adrenergic (ß2-subunit), cholinergic (M3-muscarinic) and dopaminergic (D2) receptors in the gut. Blots in (**A, C, E** and **G**) show 3 representative examples per exposure group for the protein levels of adrenergic (ß2-subunit) (B2-Adren.), cholinergic (M3-muscarinic) (M3) and dopaminergic (D2) (Dopa D2R) receptors in duodenum (**A**), jejunum (**C**), ileum (**E**) and colon (**G**). The protein gene product 9.5 (PGP 9.5) was used as a panneuronal marker to determine the neuronal population. Images were quantified using FIJI as described above. Protein expression for each marker was normalized to the amount of neuronal tissue in the sample by dividing it to the values obtained for PGP9.5 expression. Bar graphs in **B, D, F** and **H** show the quantification of the original blots (n=5). Data was analyzed using a Student’s t-test (Mann-Whitney Test). *, **, *** correspond to *P*<0.05, *P*<0.01 and *P*<0.001 respectively. n.s. non-significant. Error bars represent SEM in all graphics.

**Figure 10:**
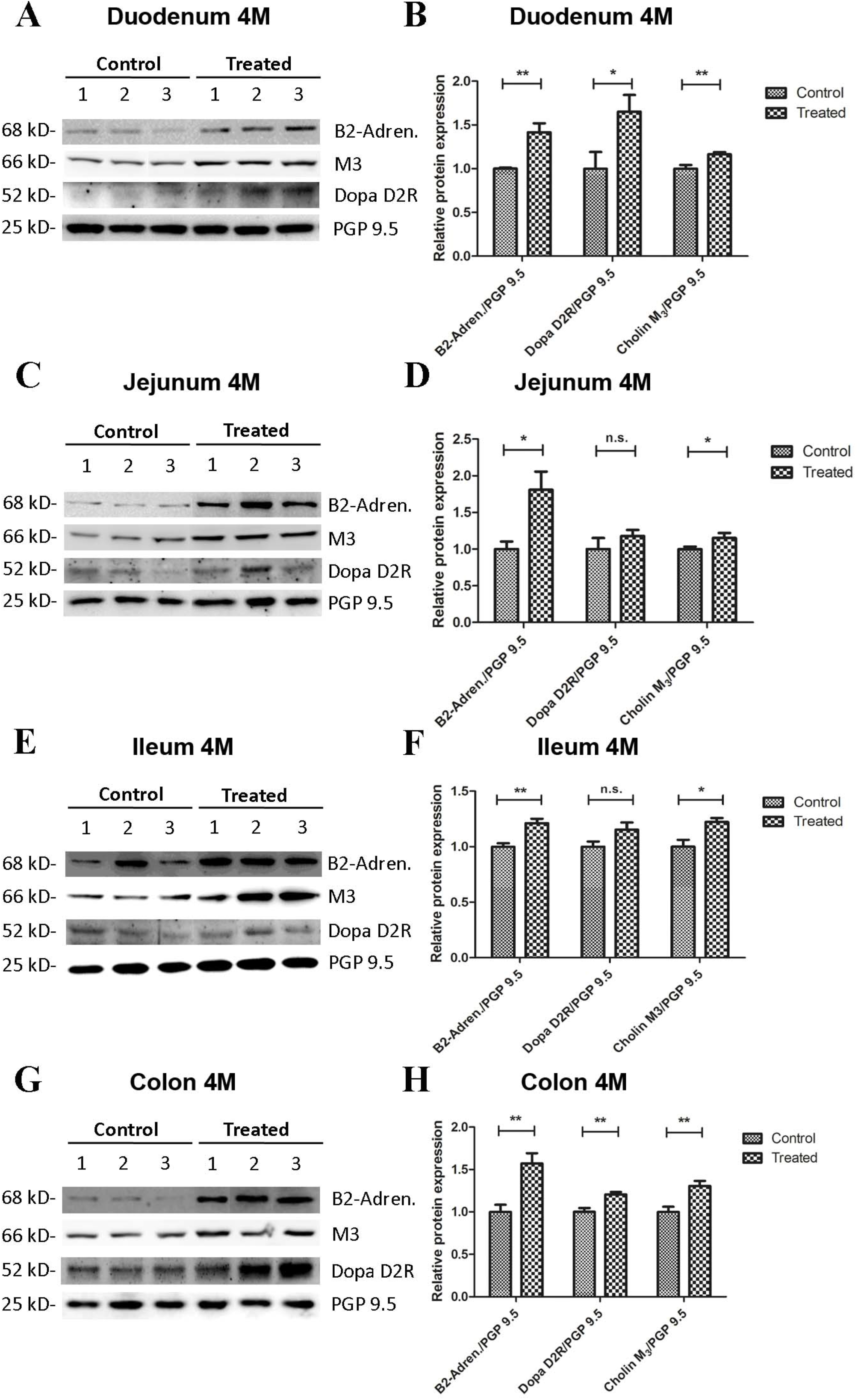
4-Month rotenone exposure induces an up regulation of the adrenergic (ß2-subunit), cholinergic (M3-muscarinic) and dopaminergic (D2) receptors in the gut. Westenr blots in (**A, C, E** and **G**) show 3 representative examples per exposure group for the protein levels of adrenergic (ß2-subunit) (B2-Adren.), cholinergic (M3-muscarinic) (M3) and dopaminergic (D2) (Dopa D2R) receptors in duodenum (**A**), jejunum (**C**), ileum (**E**) and colon (**G**). The protein gene product 9.5 (PGP 9.5) was used as a panneuronal marker to determine the total neuronal population. Images were quantified using FIJI as described in the material and methods section. Protein expression for each marker was normalized to the amount of neuronal tissue in the sample by dividing it to the values obtained for PGP9.5 expression. Bar graphs in **B, D, F** and **H** show the quantification of the original blots (n=5). Data was analyzed using a Student’s t-test (Mann-Whitney Test). *, **, *** correspond to *P*<0.05, *P*<0.01 and *P*<0.001 respectively. n.s. non-significant. Error bars represent SEM in all graphics.

### Rotenone exposure induces modifications of neuronal-specific markers in the murine intestine

In order to analyze the effect of rotenone on the enteric neurons of rotenone-exposed mice, we studied the expression of different neuronal subpopulation markers present in the duodenum, jejunum, ileum and colon from vehicle- and rotenone-exposed mice. Our results show that rotenone induces changes in all tested regions. These alterations are region and exposure dependent.

#### Exposure to rotenone does not alter PGP9.5 in the intestine of mice exposed for 2 or 4 months

To study the effect on rotenone exposure on neuronal cells we analyzed the protein expression of PGP9.5, an ubiquitin carboxy-terminal hydrolase mainly expressed in neurons and we normalized it to the expression of Glycerinaldehyd-3-phosphat-Dehydrogenase (GAPDH), which is stably expressed among the different cell populations used here as a loading control. We compared the protein expression of PGP9.5 in all intestinal regions in 2 and 4 months-rotenone- and vehicle-exposed mice. Neither analyzing all regions together (see Figure 11) or separately (see Figures 12 and 13) showed significant differences between exposure groups. We could observe minimal non-significant differences in the effect of rotenone exposure in the relative protein expression level of PGP9.5 between intestinal regions (see Figures 12 and 13). Unfortunately, we ran out of protein extract from ileum 4 months, so that no statistical analysis was possible for any of the markers tested. (See table 12)

**Figure 11:**
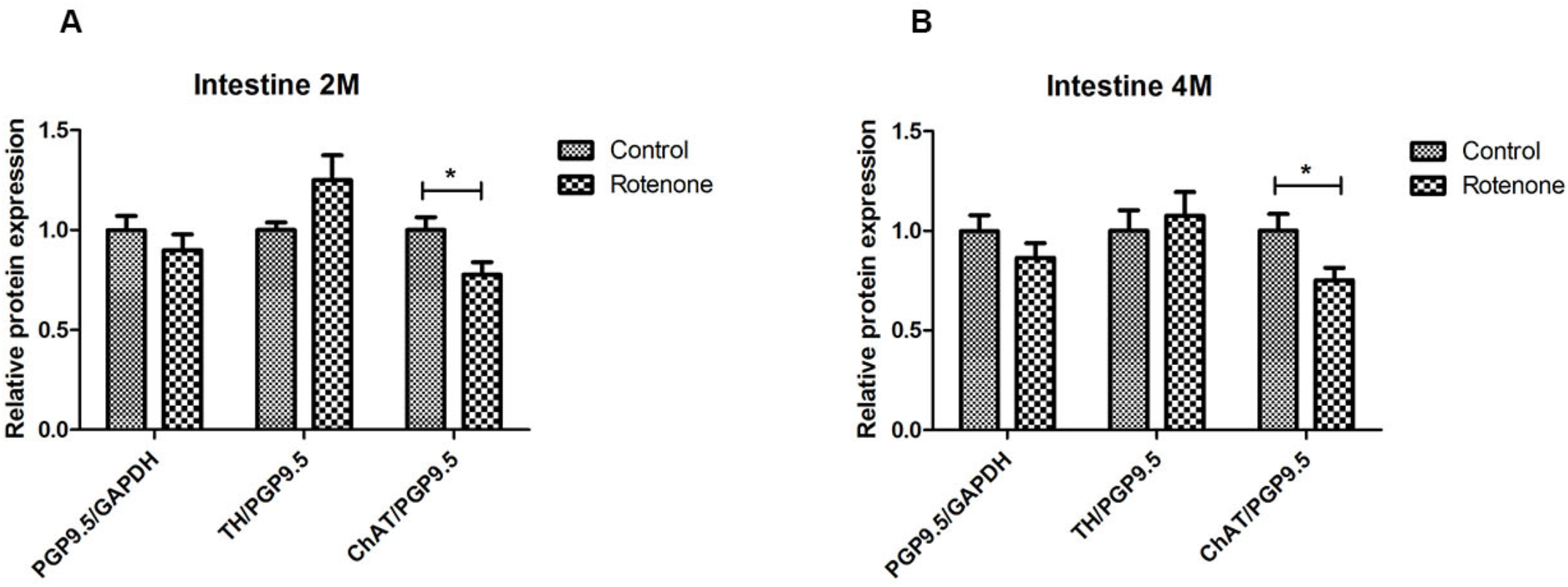
Overall effect of 2-month and 4-month rotenone exposure on the expression of ChAT, TH and PGP9.5 in the gut. Bar graph in **A** shows the protein expression levels of ChAT, TH and PGP9.5 in control and 2 months rotenone-exposed mice as the average of the values obtained for all regions. Bar graph in **B** shows the protein expression levels of ChAT, TH and PGP9.5 in 4 months control and rotenone-exposed mice as the average of the values obtained for all regions. ChAT and TH were normalized to the total neuronal tissue, determined by PGP9.5 expression levels. PGP9.5 was normalized to GAPDH. Data was analyzed using a Student’s t-test (Mann-Whitney Test). * corresponds to *P*<0.05. n.s. non-significant. Error bars represent SEM in all graphics.

**Figure 12:**
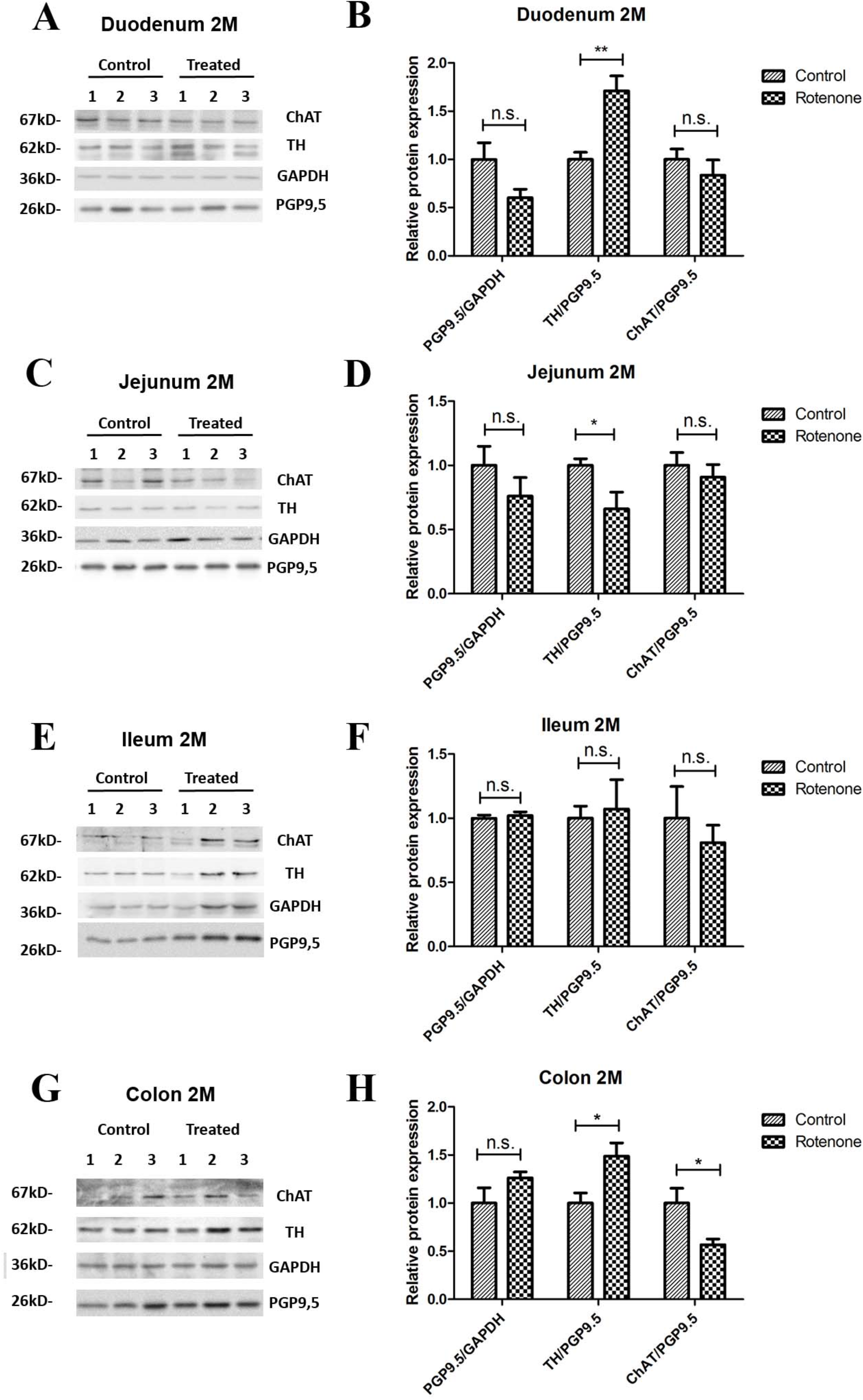
Effect of 2-month rotenone on the expression of ChAT, TH and PGP9.5 in each intestinal region. Western blots in **A, C, E** and **G** show 3 representative examples per exposure group for the protein levels of ChAT, TH, GAPDH and PGP9.5 in duodenum (**A**), jejunum (**C**), ileum (**E**) and colon (**G**). The protein gene product 9.5 (PGP 9.5) was used as a panneuronal marker to determine the ratio of enteric neuronal population within the sample. Images were quantified using FIJI as described in the material and methods section. ChAT and TH signals were normalized to the total amount of neuronal tissue in the sample by dividing it to the values obtained for PGP9.5 expression. The proportion of enteric neurons was determined by normalizing PGP9.5 signal to the total protein measured with GAPDH. Bar graphs in **B, D, F** and **H** show the quantification of the original blots (n=5). Data was analyzed using a Student’s t-test (Mann-Whitney Test). *, ** correspond to *P*<0.05 and *P*<0.01 respectively. n.s. non-significant. Error bars represent SEM in all graphics.

**Figure 13:**
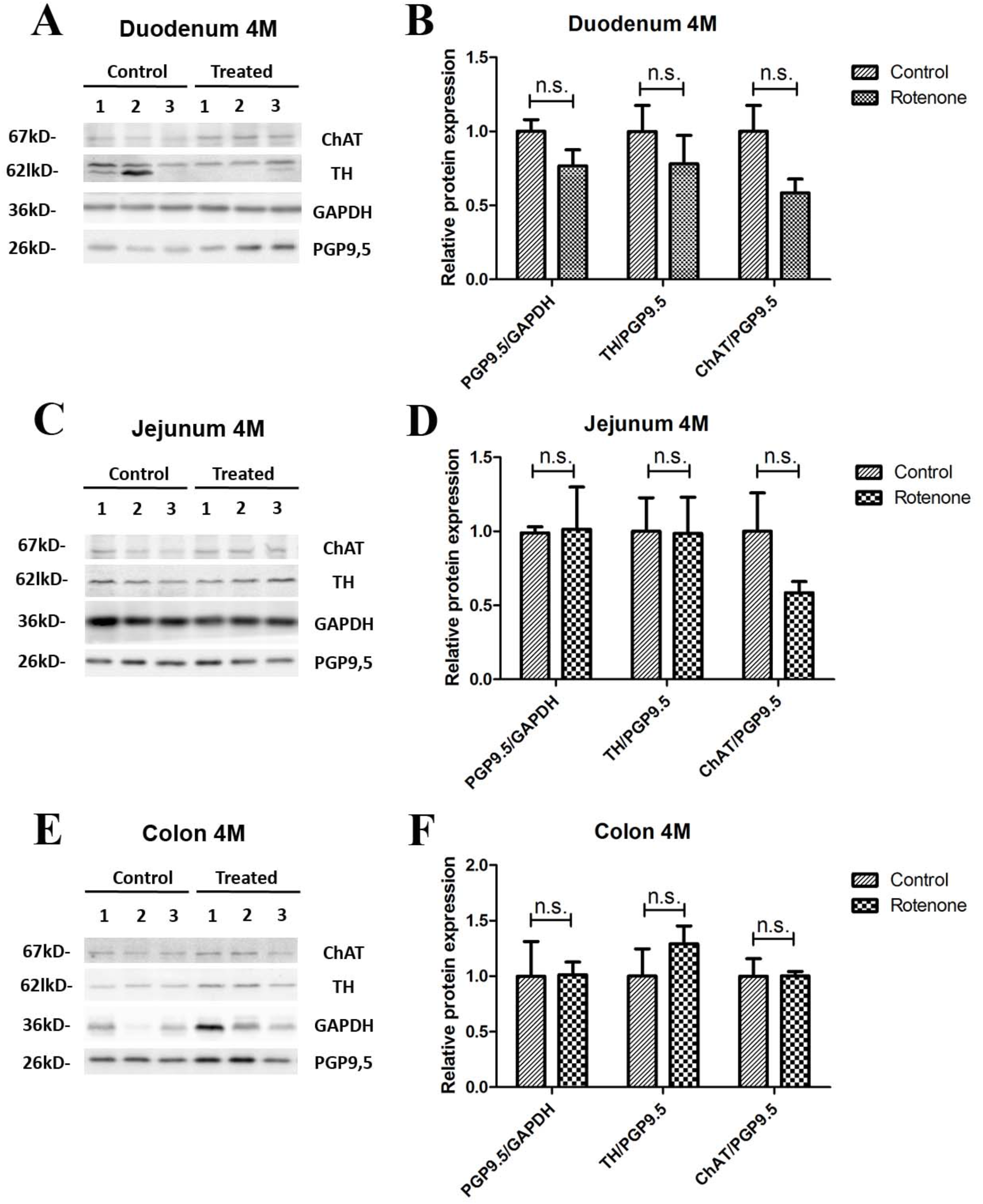
Effect of 4-month rotenone on the expression of ChAT, TH and PGP9.5 in each intestinal region. Western blots in **A, C** and **E** show 3 representative examples per exposure group for the protein levels of ChAT, TH, GAPDH and PGP9.5 in duodenum (**A**), jejunum (**C**) and colon (**E**). The protein gene product 9.5 (PGP 9.5) was used as a panneuronal marker to determine the ratio of enteric neuronal population within the sample. Images were quantified using FIJI as described in the material and methods section. ChAT and TH signals were normalized to the total amount of neuronal tissue in the sample by dividing it to the values obtained for GP9.5 expression. The proportion of enteric neurons was determined by normalizing PGP9.5 signal to the total protein measured with GAPDH. Bar graphs in **B, D** and **F** show the quantification of the original blots (n=5). Data was analyzed using a Student’s t-test. n.s. non-significant (Mann-Whitney Test). Error bars represent SEM in all graphics.

#### Exposure to rotenone for 2 and 4 months decreases enteric choline acetyltransferase levels

A reduction in the cholinergic innervation of the gut has been described in patients [17]. Therefore, we analyzed the protein expression pattern of choline acetyltransferase (ChAT), a protein expressed in cholinergic neurons, in the duodenum, jejunum, ileum and colon of control and exposed mice. When all regions were analyzed together, we observed a significant decrease in the amount of enteric ChAT expressed in rotenone-exposed mice when compared to controls after 2 months and 4 months of rotenone exposure (see Figure 11A and B). Taken separately, all intestinal regions from rotenone-exposed mice showed a decrease in ChAT expression levels when compared to controls after 2 months of exposure, which was significant only in the colon (see Figure 12). After 4 months of exposure, we observed a non-significant decrease in the amount of enteric ChAT expressed in rotenone-exposed mice when compared to controls in the duodenum and the jejunum but not in the colon (see Figure 13 and Table 13 for a summary of the results)

#### Rotenone exposure modifies tyrosine hydroxylase protein levels in the intestine of rotenone-exposed mice

We also analyzed the relative protein level of the tyrosine hydroxylase (TH/PGP9.5), implicated in the synthesis of the catecholamines dopamine, epinephrine and norepinephrine in the nervous system. We compared the protein expression of TH in all different segments from 2 and 4 months rotenone-exposed mice and controls. After 2 and 4 months of exposure, we observed no differences in the protein expression of TH in rotenone-exposed mice compared to control mice when all regions of the intestine were analyzed together (see Figure 11). However, when taken separately, TH levels were significantly higher in rotenone-exposed samples compared to control samples in duodenum and colon after two months of rotenone exposure. In the meantime, no difference was observed in the ileum and TH levels were lower in the jejunum of rotenone-exposed mice, highlighting the region-specific effect of rotenone exposure in the intestine. After 4 months of rotenone exposure we observed a non-significant increase in TH protein expression in the colon of rotenone-exposed mice compared to control mice whereas this expression was either similar in the jejunum or lower in the duodenum of rotenone-exposed mice compared to control mice. (see Figures 12 and 13 and Table 14).

#### Changes in the level of α_4_β_2_* nicotinic acetylcholine receptors in the gastrointestinal tract are not detectable by PET after 2 or 4 months exposure to rotenone

To evaluate the impact of exposure to rotenone on the cholinergic innervation of the gut, we performed *in vivo* PET imaging. We selected a clinically used radiotracer, (−)-[^18^F]Flubatine, selective of the α_4_β_2_* nicotinic acetylcholine receptor subtype. Two regions were selected for receptor density evaluation: the jejunum and the caecum. After 2 months of exposure to rotenone no alteration of α_4_β_2_* nicotinic acetylcholine receptors density could be detected with PET imaging in the jejunum as well as in the caecum region when compared to controls (see Figure 14 A-C). After 4 months of exposure to rotenone we observed a non-significant decrease in the density of enteric α_4_β_2_* nicotinic acetylcholine receptors in the jejunum and the caecum when compared to controls (see Figure 14 B-D).

**Figure 14:**
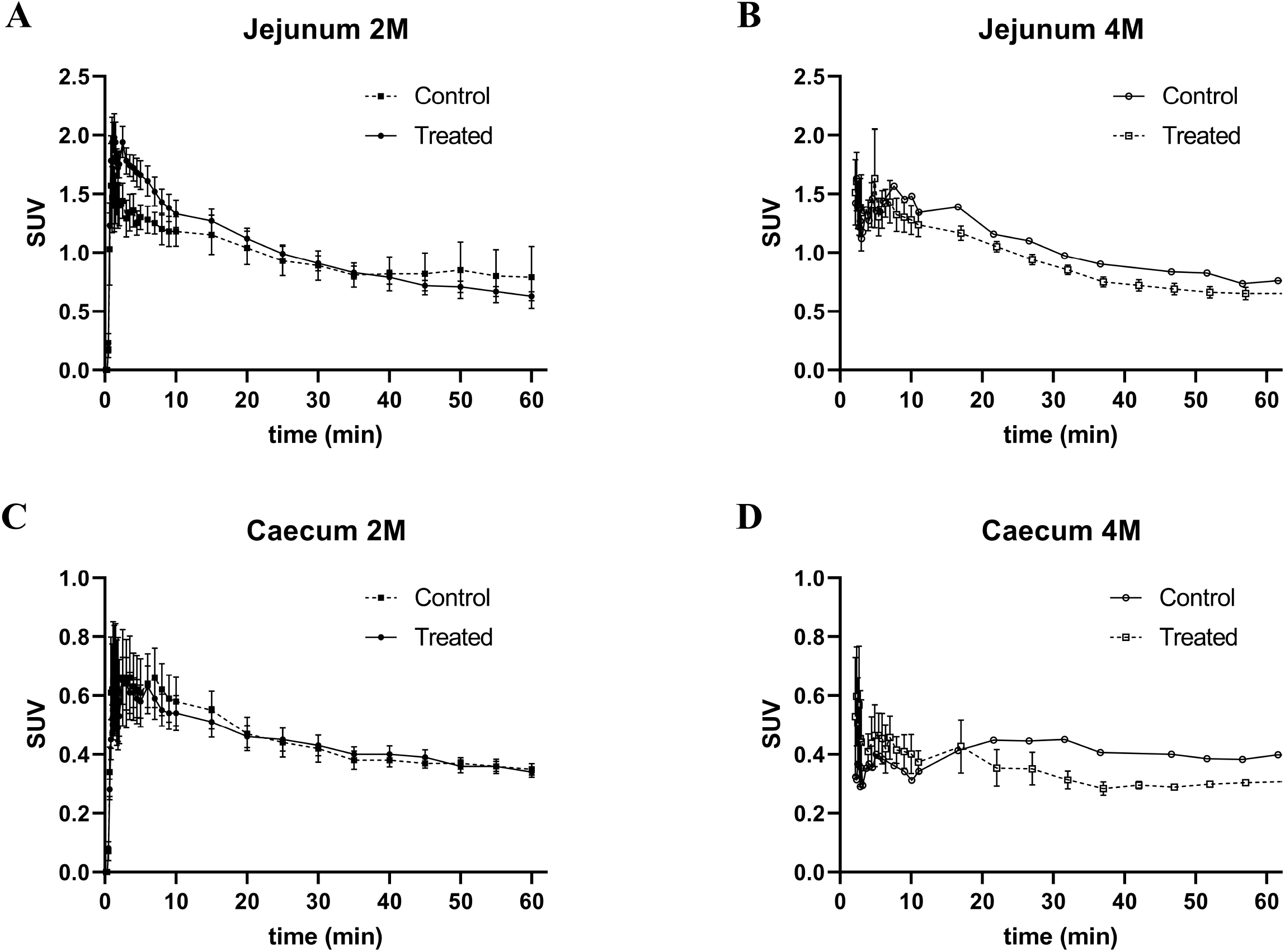
(−)-[^18^F]Flubatine-PET imaging of enteric α_4_β_2_* nicotinic acetylcholine receptor density after 4 months exposure to rotenone. Time-activity curves of the jejunum after exposure to rotenone of 2 months (2M) (**A**) or 4 months (4M) (**B)**; time-activity curves of the caecum after exposure to rotenone of 2 months (2M) (**C**) or 4 months (4M) (**D**); after injection of (−)-[^18^F]Flubatine in C57/BL6J mice control or treated. Error bars represent SEM.

## Discussion

Non-motor symptoms in Parkinson’s disease are widely spread among patients. The prevalence varies depending on the type of symptom and, unlike motor symptoms, they are not always observed in PD patients [18–24]. According to Braak’s pathological staging, most of the impaired nervous structures giving rise to non-motor symptoms (e.g. enteric, sympathetic or parasympathetic systems) are affected in very early stages of the disease and show PD-pathology in the form of LB and LN [1]. Specifically, gastrointestinal alterations can appear up to 15 years before the appearance of the motor symptoms in PD patients. In this study we focused on the analysis of gastrointestinal symptoms in a toxic model of Parkinson’s disease based on the oral administration of rotenone. In previous studies we have shown that the exposure to oral administered rotenone induces PD-like changes in the ENS and triggers the progression of PD throughout the nervous system until it reaches the substantia nigra [25, 26]. Complementarily, the results of the present study suggest that the gastrointestinal alterations present in PD patients could also be explained by such an exposure to environmental toxins due to the modification of the neuronal population, the impairment of the vagal and sympathetic innervation of the ENS and the subsequent up-regulation in the expression of cholinergic, adrenergic and dopaminergic receptors within the intestine in a region-, age- and exposure-dependent manner. Interestingly, the latter changes appear already at early time-points after rotenone administration (2 months) before the onset of motor symptoms (that in this animal model occur after 3 months of exposure), thus, mimicking the progression pattern observed in PD patients. The oral exposure to rotenone in this model mimics a realistic exposure to pesticide as it occurs in the general population and would also explain that the intestinal symptoms are the first to appear.

In two recent studies, it was shown that rotenone exposure reduces the sympathetic noradrenergic [13] and the vagal cholinergic [14] innervation of the gut. In order to determine whether enteric neurotransmitters did also play a role in these alterations, we analyzed the differences in the total enteric neuronal population and its subpopulations between control and rotenone-exposed mice. Although there was a general decrease in the expression of PGP9.5, ChAT and TH in rotenone-exposed mice (suggesting a general neuronal loss) when analyzed by intestinal regions, this decrease was not significant in most regions. Moreover, the changes observed in the different analyzed regions did not remain constant with time, even under exposure, suggesting the presence of neuronal plasticity in response to pesticide exposure. This could also provide an explanation for the contradictory results between regions and within the same region between different exposure times. In this regard, it was recently shown that the enteric nervous system is able to regenerate through neurogenesis [27, 28], a process already described in other degenerative processes of the gut. These results would also agree with a recent study published by Annerino and colleagues [29]. In this study they did not observe a neuronal loss in the myenteric ganglia of PD patients when compared to controls.

Overall these results support the hypothesis that the sympathetic and cholinergic innervation of the gut plays a main role in the pathophysiology of the functional gastrointestinal alterations and that the role of the intrinsic neurotransmitters is, if something, marginal and transitory. How the extrinsic innervation of the gut was able to alter the expression of TH in the gut remains unclear. We speculate that the activation of postsynaptic enteric neurons by ACh and NA in the ENS leads to the activation of dopaminergic neurons and in the absence of this input there is an up-regulation in the expression of enteric dopaminergic receptors, but it needs further clarification.

The early appearance, the wide consistency across time and intestinal regions of this up-regulation and the accessibility to obtain biopsies (specially for the colon) make these biomarkers, in combination with the diagnosis of other premotor symptoms such as idiopathic REM sleep behavior disorder, ideal for the early detection of PD. Therefore, we analyzed if there were variations in the expression of the α4β2* nicotinic cholinergic receptor using (−)-[18F]Flubatine as a radiotracer. (−)-[18F]Flubatine has been previously used in the clinic as a biomarker for Alzheimer’s disease [30] and is not eliminated through the intestine, an important pharmacokinetic characteristic when investigating expression levels in the gut. The results only showed a non-significant reduction in the PET signal intensity. In this regard, it is important to note that the metabolism of (−)-[^18^F]Flubatine is higher in mice than human [31] and, together with the peristaltic movement, could be confounding factors for the PET data evaluation. Additionally, the PET signal cannot be normalised to the amount of neurons present in that region as was done with the western-blot results. Therefore, it could be possible that the signal observed is the result of an up regulated receptor expression in a diminished neuronal population. Finally, it has been shown that α4β2* nicotinic cholinergic receptors expressed in the brain have a paradoxical effect to stimulation with nicotine, as they are up regulated by it [32, 33] . It is therefore possible that a reduction in the cholinergic input could lead to a down regulation of this receptor instead of an up regulation. Overall, these findings may be in favour of a cholinergic denervation as described in patients [31], but further investigation using radiotracers targeting other receptors would be needed in order to discriminate properly. These results suggest that using a suitable radiotracer, or a combination of them, together with other diagnostic techniques (i.e. EEG for REM-sleep disorders or high-speed MRI for gastric motility alterations) could increase the sensibility and specificity in the early diagnosis of PD, which would allow for an earlier therapeutic intervention and a better outcome.

## Supporting information

Quantification code MatLab

## Authors Contribution

FP-M designed the study. YD exposed the mice. UR and FP-M designed the contractility experiments. RM, YD and FP-M performed the contractility experiments. KB wrote the Matlab quantification program. UR, GS, KB and FP-M analyzed the contractility results. AS, QS and FP-M performed the western-blots. CV-B, AS, QS, RF and FP-M analyzed the western-blot results. PB, MT, MK, DG performed the PET and analyzed the PET data. MD, KB, RF and FP-M wrote and critically corrected the manuscript.

## Acknowledgements

we would like to thank Prof. Marianne Patt and Prof. Osama Sabri from the Department of Nuclear Medicine of the Leipzig University for kindly providing us with the PET-radiotracer used in this study.

