## Supplementary Figures for "Physiological and pathological changes in the enteric nervous system of rotenone-exposed mice as early biomarkers for Parkinson’s disease"

Supp. Fig. 1

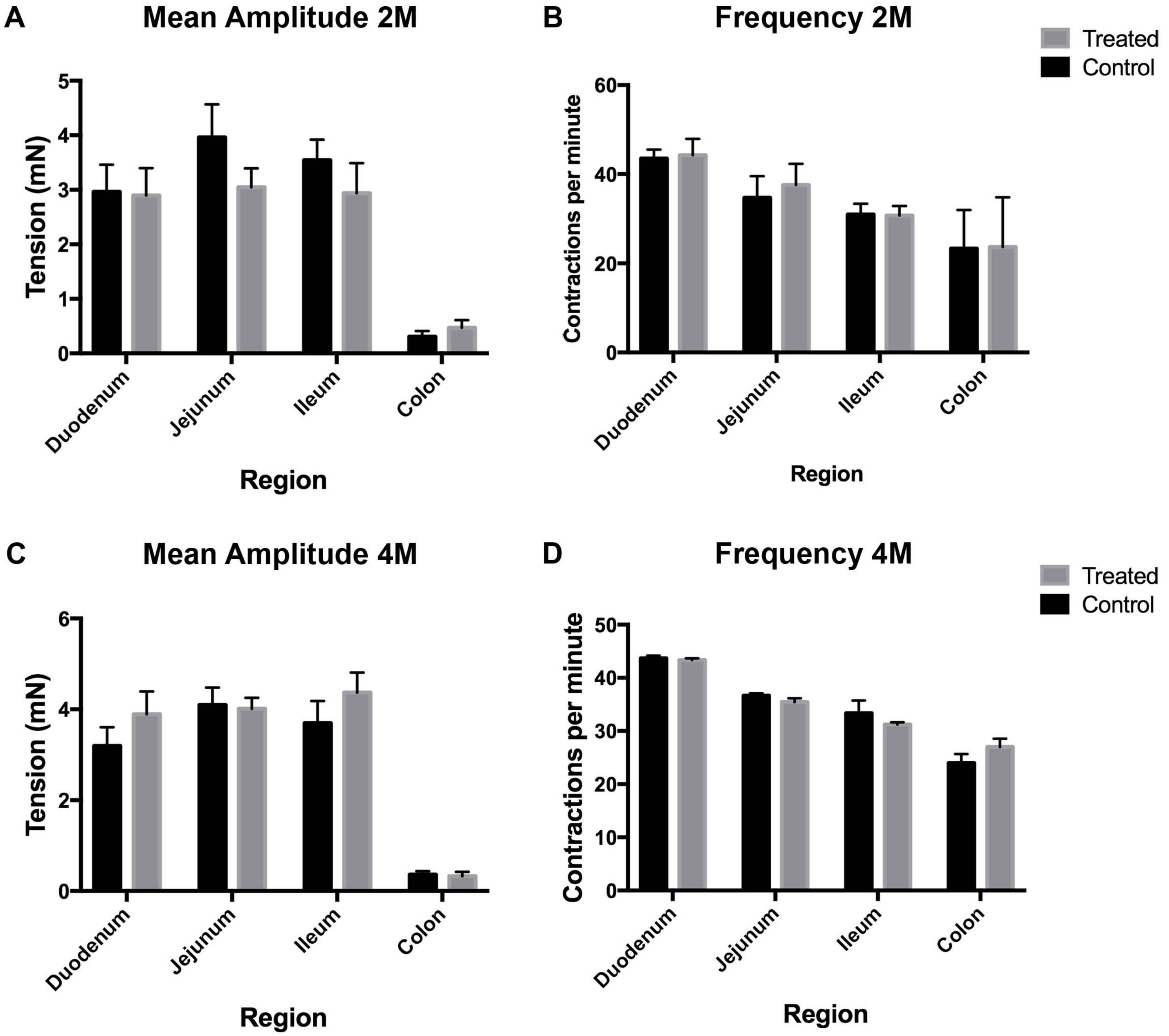
A
Mean Amplitude 2M
Tension (mN)
0
1
2
3
4
5
Duodenum
Jejunum
Ileum
Colon
Region
B
Frequency 2M
Contractions per minute
0
20
40
60
Duodenum
Jejunum
Ileum
Colon
Region
Treated
Control
C
Mean Amplitude 4M
Tension (mN)
0
2
4
6
Duodenum
Jejunum
Ileum
Colon
Region
D
Frequency 4M
Contractions per minute
0
10
20
30
40
50
Duodenum
Jejunum
Ileum
Colon
Region
Treated
Control

Supp. Fig. 2

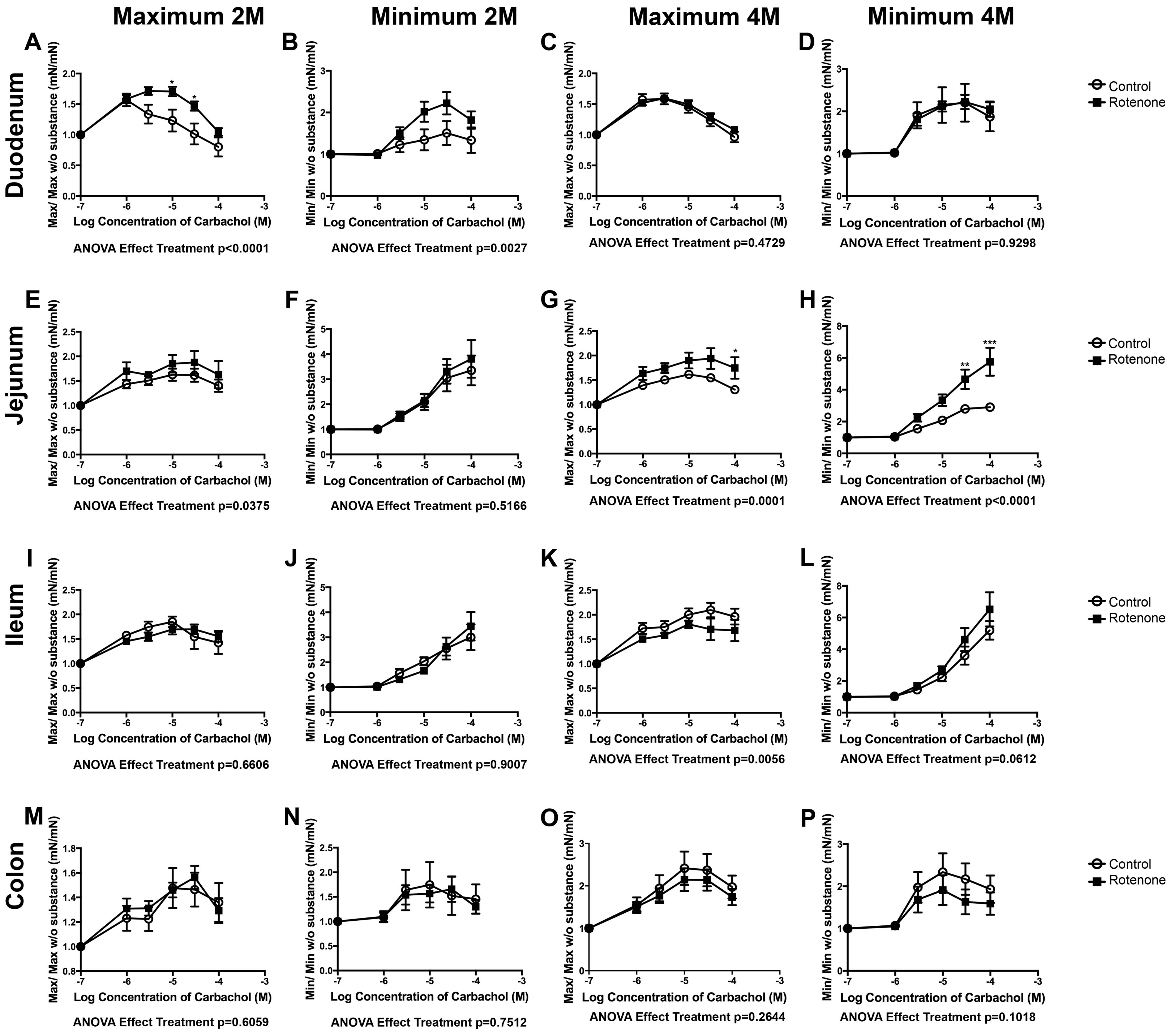

Maximum 2M
Minimum 2M
Maximum 4M
Minimum 4M
Duodenum
Jejunum
Ileum
Colon
A
B
C
D
E
F
G
H
I
J
K
L
M
N
O
P
Max/ Max w/o substance (mN/mN)
Min/ Min w/o substance (mN/mN)
Log Concentration of Carbachol (M)
Control
Rotenone
ANOVA Effect Treatment p<0.0001
ANOVA Effect Treatment p=0.0027
ANOVA Effect Treatment p=0.4729
ANOVA Effect Treatment p=0.9298
ANOVA Effect Treatment p=0.0375
ANOVA Effect Treatment p=0.5166
ANOVA Effect Treatment p=0.0001
ANOVA Effect Treatment p<0.0001
ANOVA Effect Treatment p=0.6606
ANOVA Effect Treatment p=0.9007
ANOVA Effect Treatment p=0.0056
ANOVA Effect Treatment p=0.0612
ANOVA Effect Treatment p=0.6059
ANOVA Effect Treatment p=0.7512
ANOVA Effect Treatment p=0.2644
ANOVA Effect Treatment p=0.1018

Supp. Fig. 3

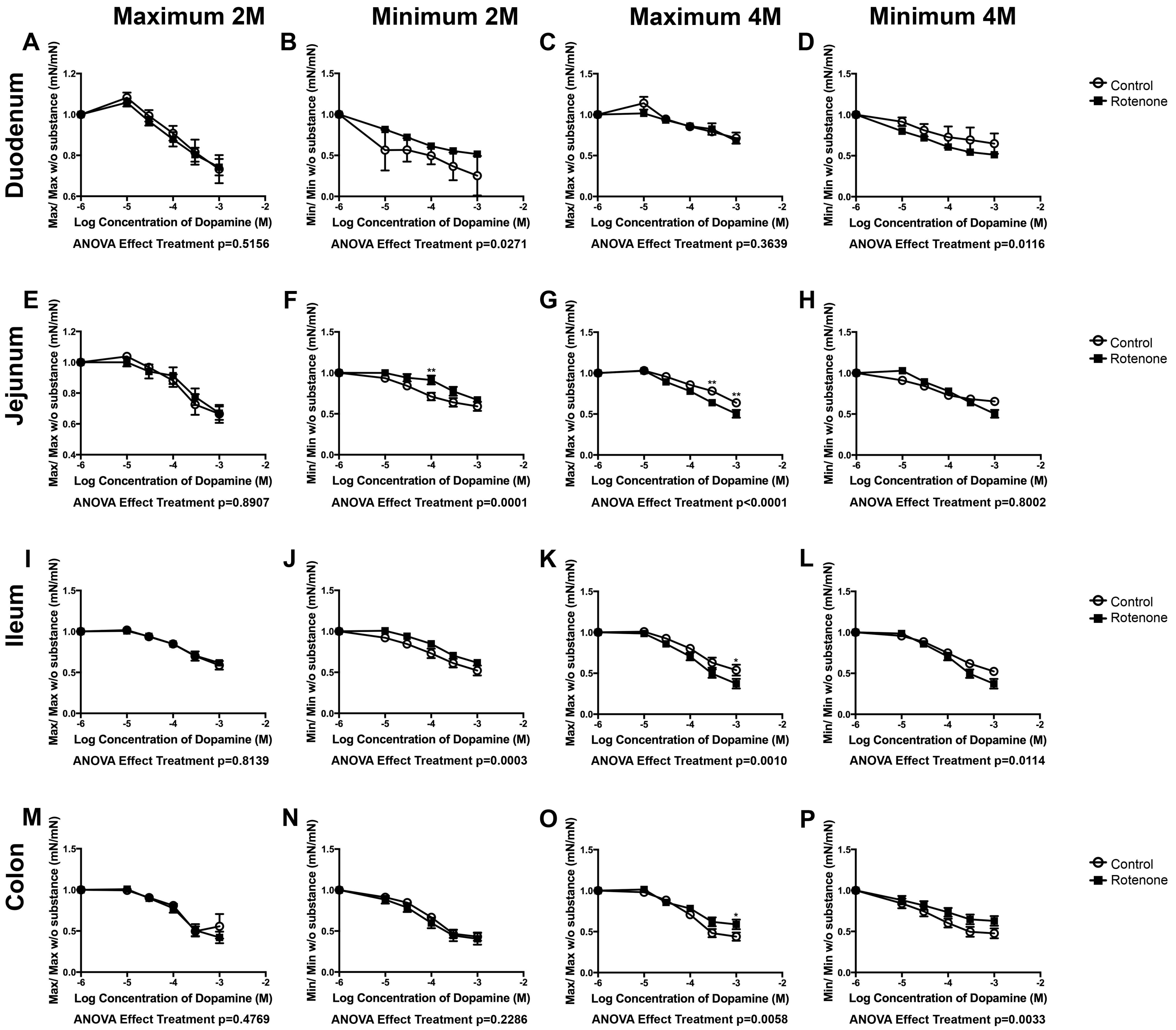
Maximum 2M
Minimum 2M
Maximum 4M
Minimum 4M
Duodenum
Jejunum
Ileum
Colon
A
B
C
D
E
F
G
H
I
J
K
L
M
N
O
P
Control
Rotenone
Max/ Max w/o substance (mN/mN)
Min/ Min w/o substance (mN/mN)
Log Concentration of Dopamine (M)
ANOVA Effect Treatment p=0.5156
ANOVA Effect Treatment p=0.0271
ANOVA Effect Treatment p=0.3639
ANOVA Effect Treatment p=0.0116
ANOVA Effect Treatment p=0.8907
ANOVA Effect Treatment p=0.0001
ANOVA Effect Treatment p<0.0001
ANOVA Effect Treatment p=0.8002
ANOVA Effect Treatment p=0.8139
ANOVA Effect Treatment p=0.0003
ANOVA Effect Treatment p=0.0010
ANOVA Effect Treatment p=0.0114
ANOVA Effect Treatment p=0.4769
ANOVA Effect Treatment p=0.2286
ANOVA Effect Treatment p=0.0058
ANOVA Effect Treatment p=0.0033

Supp. Fig. 4

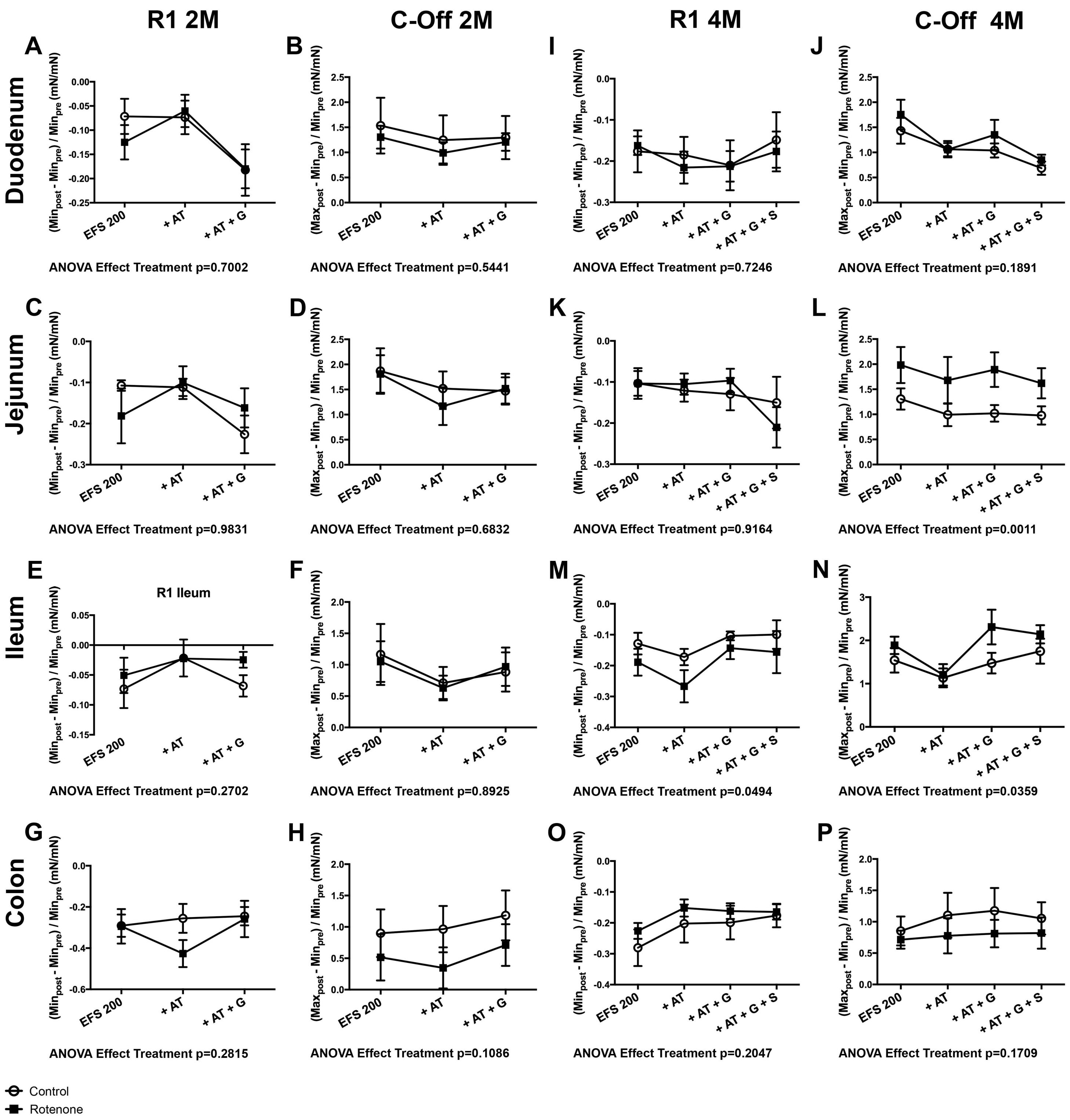
R1 2M
C-Off 2M
R1 4M
C-Off 4M
Duodenum
Jejunum
Ileum
Colon
A
B
I
J
C
D
K
L
E
F
M
N
G
H
O
P
R1 Ileum
EFS 200
+ AT
+ AT + G
+ AT + G + S
ANOVA Effect Treatment p=0.7002
ANOVA Effect Treatment p=0.5441
ANOVA Effect Treatment p=0.7246
ANOVA Effect Treatment p=0.1891
ANOVA Effect Treatment p=0.9831
ANOVA Effect Treatment p=0.6832
ANOVA Effect Treatment p=0.9164
ANOVA Effect Treatment p=0.0011
ANOVA Effect Treatment p=0.2702
ANOVA Effect Treatment p=0.8925
ANOVA Effect Treatment p=0.0494
ANOVA Effect Treatment p=0.0359
ANOVA Effect Treatment p=0.2815
ANOVA Effect Treatment p=0.1086
ANOVA Effect Treatment p=0.2047
ANOVA Effect Treatment p=0.1709
Control
Rotenone

Supp. Fig. 5

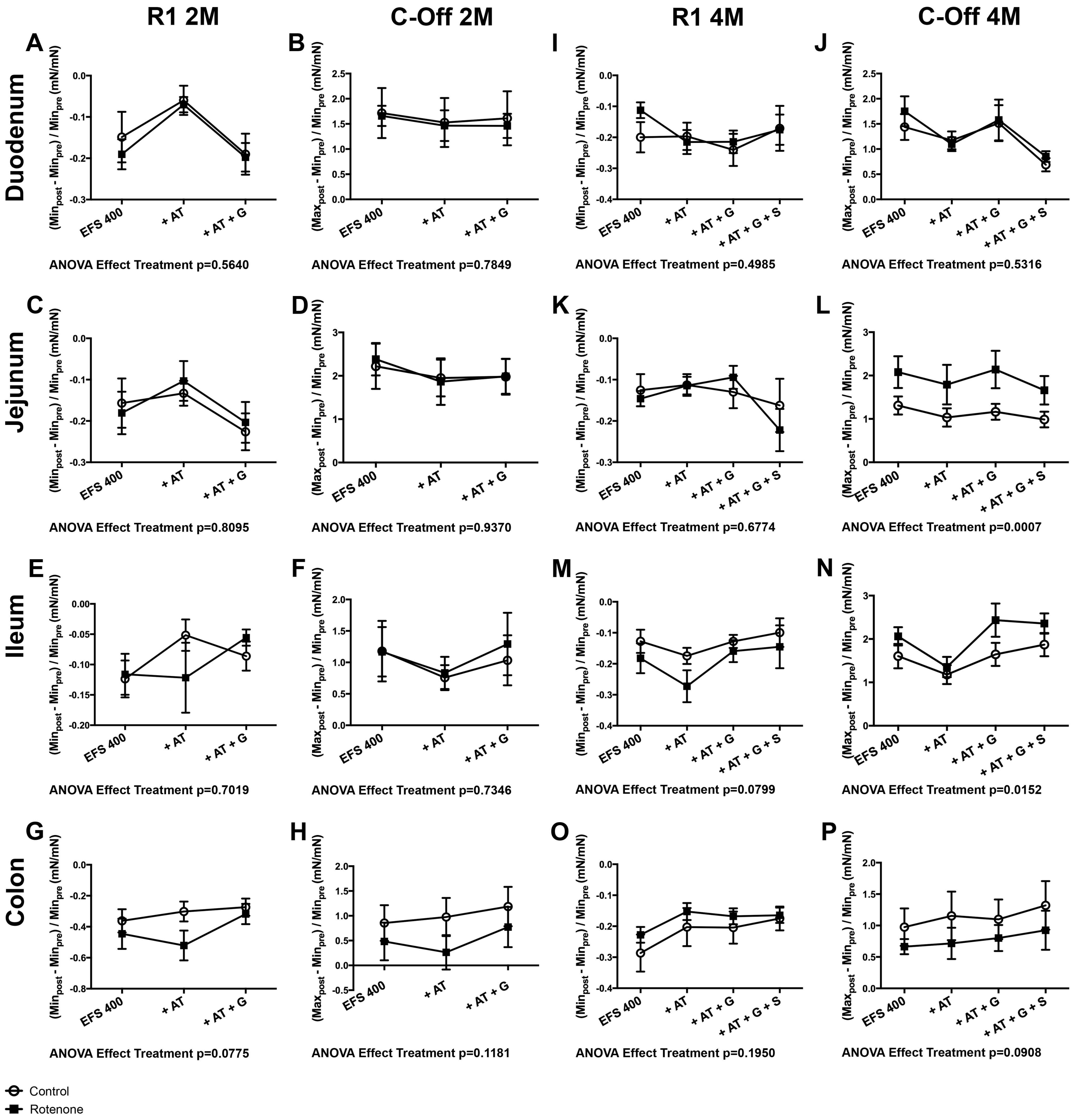
R1 2M
C-Off 2M
R1 4M
C-Off 4M
Duodenum
Jejunum
Ileum
Colon
A
B
I
J
C
D
K
L
E
F
M
N
G
H
O
P
(Min_post - Min_pre) / Min_pre (mN/mN)
(Max_post - Min_pre) / Min_pre (mN/mN)
EFS 400
+ AT
+ AT + G
+ AT + G + S
ANOVA Effect Treatment p=0.5640
ANOVA Effect Treatment p=0.7849
ANOVA Effect Treatment p=0.4985
ANOVA Effect Treatment p=0.5316
ANOVA Effect Treatment p=0.8095
ANOVA Effect Treatment p=0.9370
ANOVA Effect Treatment p=0.6774
ANOVA Effect Treatment p=0.0007
ANOVA Effect Treatment p=0.7019
ANOVA Effect Treatment p=0.7346
ANOVA Effect Treatment p=0.0799
ANOVA Effect Treatment p=0.0152
ANOVA Effect Treatment p=0.0775
ANOVA Effect Treatment p=0.1181
ANOVA Effect Treatment p=0.1950
ANOVA Effect Treatment p=0.0908
Control
Rotenone

**Supplementary figure 1: Mean Amplitude and Frequency of contractions in the regions of the intestine after 2 and 4 months of exposure to rotenone.** Bar graphics in **A** to **D** show the comparison of the Mean Amplitude (**A** and **C**) and the Frequency of contraction (**B** and **D**) after 2 months (**A** and **B**) and 4 months (**C** and **D**) of rotenone exposure. No significant difference was observed in any of the parameters analyzed. Error bars represent SEM.

**Supplementary figure 2: Comparison between the concentration curves of carbachol and its effects on the different regions of the intestine between control and rotenone exposed mice.** Graphics in A to P show the effect of different concentrations of carbachol on the maximal exerted contractility force (Max) first and third columns and on the intestinal muscle tone (Min) second and fourth columns in two months (2M) and four months (4M) vehicle (empty circle) and rotenone exposed (black square) mice. *, *** correspond to $P<0.05$ and $P<0.001$ respectively. Error bars represent SEM.

**Supplementary figure 3: Comparison between the concentration curves of dopamine and its effects on the different regions of the intestine between control and rotenone exposed mice.** Graphics in A to P show the effect of different concentrations of dopamine on the maximal exerted contractility force (Max) first and third columns and on the intestinal muscle tone (Min) second and fourth columns in two months (2M) and four months (4M) vehicle (empty circle) and rotenone exposed (black square) mice. ** correspond to $P<0.01$. Error bars represent SEM.

**Supplementary figure 4: Comparison of the effect of a 200 µs electric field stimulation (EFS 200) on the different intestinal regions after 2 and 4 months of rotenone exposure in mice.** Graphics in A to P show variations in the R1 (first and third columns) and C-Off (second and fourth columns) responses to the EFS 200 alone and after the addition of AT alone, AT and G or AT, G and S (serotonin only after 4 months of exposure). The overall response was compared between control and rotenone exposed mice with a 2-way ANOVA. Significance is written under each graphic.

**Supplementary figure 5: Comparison of the effect of a 400 µs electric field stimulation (EFS 400) on the different intestinal regions after 2 and 4 months of rotenone exposure in mice.** Graphics in A to P show variations in the in the R1 (first and third columns) and C-Off (second and fourth columns) responses to the EFS 400 alone and after the addition of AT alone, AT and G or AT, G and S (serotonin only after 4 months of exposure). The overall response was compared between control and rotenone exposed mice with a 2-way ANOVA. Significance is written under each graphic.

### Table 1

| Carbachol | | 2-way ANOVA Treatment p value<br>p mean amplitude | p maximum | p minimum |
|---|---|---|---|---|
| Duodenum | 2M | **<0,001** | **<0,001** | **0,003** |
| | 4M | 0,263 | 0,473 | 0,930 |
| Jejunum | 2M | **<0,001** | **0,038** | 0,517 |
| | 4M | **<0,001** | **<0,001** | **<0,001** |
| Ileum | 2M | 0,009 | 0,660 | 0,901 |
| | 4M | 0,288 | **0,006** | 0,061 |
| Colon | 2M | 0,085 | 0,606 | 0,751 |
| | 4M | **0,043** | 0,264 | 0,102 |

### Table 2

| Dopamine | | 2-way ANOVA Treatment p value<br>p mean amplitude | p maximum | p minimum |
|---|---|---|---|---|
| Duodenum | 2M | 0,395 | 0,516 | **0,027** |
| | 4M | **0,022** | 0,364 | **0,012** |
| Jejunum | 2M | 0,689 | 0,891 | **<0,001** |
| | 4M | **0,012** | **<0,001** | 0,800 |
| Ileum | 2M | 0,890 | 0,814 | **<0,001** |
| | 4M | **0,009** | **0,001** | **0,011** |
| Colon | 2M | 0,186 | 0,477 | 0,229 |
| | 4M | 0,286 | **0,006** | **0,003** |

### Table 3

| R1 AT | | Control | | p | Rotenone | | p | p Control vs Rotenone |
|---|---|---|---|---|---|---|---|---|
| | | Mean | SEM | | Mean | SEM | | |
| Duodenum | 2M EFS 200 | 315,191 | 175,939 | 0,249 | 116,545 | 85,781 | 0,851 | 0,333 |
| | 2M EFS 400 | 41,398 | 20,512 | **0,017** | 44,004 | 18,337 | **0,012** | 0,926 |
| | 4M EFS 200 | 146,393 | 45,084 | 0,290 | 153,464 | 41,517 | 0,216 | 0,148 |
| | 4M EFS 400 | 168,833 | 69,876 | 0,311 | 215,126 | 92,574 | 0,232 | 0,342 |
| Jejunum | 2M EFS 200 | 118,449 | 33,240 | 0,591 | 72,171 | 42,528 | 0,528 | 0,397 |
| | 2M EFS 400 | 155,139 | 67,374 | 0,432 | 66,621 | 31,924 | 0,349 | 0,272 |
| | 4M EFS 200 | 201,091 | 49,656 | 0,059 | 81,788 | 18,877 | 0,349 | **0,018** |
| | 4M EFS 400 | 134,984 | 42,752 | 0,425 | 83,759 | 13,506 | 0,247 | 0,499 |
| Ileum | 2M EFS 200 | 341,479 | 280,242 | 0,409 | 122,493 | 46,901 | 0,642 | 0,459 |
| | 2M EFS 400 | 30,042 | 21,168 | **0,008** | 136,076 | 36,440 | 0,346 | **0,031** |
| | 4M EFS 200 | 257,809 | 102,348 | 0,121 | 185,114 | 57,208 | 0,156 | 0,116 |
| | 4M EFS 400 | 59,825 | 6,443 | **0,033** | 155,883 | 31,187 | 0,092 | 0,908 |
| Colon | 2M EFS 200 | 80,848 | 15,441 | 0,243 | 230,193 | 98,750 | 0,217 | 0,166 |
| | 2M EFS 400 | 59,825 | 6,443 | 0,283 | 122,885 | 16,826 | 0,204 | 0,107 |
| | 4M EFS 200 | 63,827 | 14,303 | **0,022** | 70,743 | 10,085 | **0,010** | 0,698 |
| | 4M EFS 400 | 63,568 | 14,458 | **0,022** | 70,603 | 9,899 | **0,009** | 0,693 |

### Table 4

| C-Off AT | | Control | | p | Rotenone | | p | p Control vs Rotenone |
|---|---|---|---|---|---|---|---|---|
| | | Mean | SEM | | Mean | SEM | | |
| Duodenum | 2M EFS 200 | 79,173 | 14,143 | 0,171 | 73,393 | 11,249 | **0,040** | 0,756 |
| | 2M EFS 400 | 95,322 | 20,113 | 0,821 | 90,426 | 18,887 | 0,623 | 0,863 |
| | 4M EFS 200 | 81,353 | 9,268 | **0,049** | 65,884 | 6,538 | **<0,001** | 0,185 |
| | 4M EFS 400 | 88,081 | 8,791 | 0,169 | 68,931 | 6,566 | **<0,001** | 0,097 |
| Jejunum | 2M EFS 200 | 95,973 | 15,359 | 0,799 | 78,510 | 32,023 | 0,517 | 0,634 |
| | 2M EFS 400 | 98,213 | 15,010 | 0,908 | 75,032 | 14,189 | 0,109 | 0,288 |
| | 4M EFS 200 | 68,503 | 10,650 | **0,009** | 79,165 | 10,480 | 0,064 | 0,486 |
| | 4M EFS 400 | 74,035 | 7,839 | **0,004** | 82,448 | 10,150 | 0,103 | 0,521 |
| Ileum | 2M EFS 200 | 73,371 | 25,338 | 0,318 | 65,949 | 12,992 | **0,026** | 0,8 |
| | 2M EFS 400 | 80,838 | 20,061 | 0,362 | 76,370 | 10,713 | 0,052 | 0,848 |
| | 4M EFS 200 | 81,073 | 21,466 | 0,363 | 59,825 | 6,443 | **<0,001** | 0,334 |
| | 4M EFS 400 | 76,077 | 14,726 | 0,104 | 63,085 | 6,077 | **<0,001** | 0,408 |
| Colon | 2M EFS 200 | 99,826 | 36,553 | 0,996 | 27,022 | 11,878 | **<0,001** | 0,087 |
| | 2M EFS 400 | 94,823 | 47,659 | 0,916 | -16,350 | 27,606 | **0,002** | 0,08 |
| | 4M EFS 200 | 109,166 | 16,862 | 0,594 | 97,017 | 13,881 | 0,833 | 0,586 |
| | 4M EFS 400 | 104,363 | 11,605 | 0,712 | 97,498 | 16,565 | 0,882 | 0,739 |

### Table 5

| R1 AT + G | | Control AT | | Control AT + G | | p | Rotenone AT | | Rotenone AT + G | | p | p Control vs Rotenone |
|---|---|---|---|---|---|---|---|---|---|---|---|---|
| | | Mean | SEM | Mean | SEM | | Mean | SEM | Mean | SEM | | |
| Duodenum | 2M EFS 200 | 315,191 | 175,939 | 813,131 | 426,648 | 0,315 | 116,545 | 85,781 | 217,024 | 115,050 | 0,500 | 0,216 |
| | 2M EFS 400 | 41,398 | 20,512 | 162,965 | 57,991 | 0,076 | 44,004 | 18,337 | 108,959 | 19,453 | **0,035** | 0,398 |
| | 4M EFS 200 | 146,393 | 45,084 | 129,178 | 24,202 | 0,742 | 153,464 | 41,517 | 181,382 | 55,165 | 0,691 | 0,169 |
| | 4M EFS 400 | 168,833 | 69,876 | 267,884 | 148,191 | 0,555 | 215,126 | 92,574 | 218,659 | 89,956 | 0,979 | 0,25 |
| Jejunum | 2M EFS 200 | 118,449 | 33,240 | 227,785 | 52,248 | 0,108 | 72,171 | 42,528 | 118,189 | 49,128 | 0,495 | 0,157 |
| | 2M EFS 400 | 155,139 | 67,374 | 227,716 | 76,153 | 0,492 | 68,621 | 31,924 | 124,575 | 24,811 | 0,196 | 0,227 |
| | 4M EFS 200 | 201,091 | 49,656 | 300,511 | 132,088 | 0,491 | 81,788 | 18,877 | 83,871 | 22,815 | 0,945 | 0,083 |
| | 4M EFS 400 | 134,984 | 42,752 | 128,653 | 23,275 | 0,898 | 83,759 | 13,506 | 78,240 | 22,770 | 0,838 | 0,746 |
| Ileum | 2M EFS 200 | 341,479 | 280,242 | 565,512 | 454,462 | 0,684 | 122,493 | 46,901 | 82,262 | 44,370 | 0,547 | 0,315 |
| | 2M EFS 400 | 30,042 | 21,168 | 76,008 | 27,997 | 0,219 | 136,076 | 36,440 | 68,049 | 36,293 | 0,215 | 0,866 |
| | 4M EFS 200 | 257,809 | 102,348 | 202,297 | 101,587 | 0,706 | 185,114 | 57,208 | 128,629 | 59,287 | 0,503 | 0,125 |
| | 4M EFS 400 | 172,554 | 33,006 | 136,134 | 33,290 | 0,450 | 155,883 | 31,187 | 104,455 | 29,719 | 0,250 | 0,61 |
| Colon | 2M EFS 200 | 80,848 | 15,441 | 83,618 | 6,007 | 0,871 | 230,193 | 98,750 | 70,959 | 24,153 | 0,148 | 0,622 |
| | 2M EFS 400 | 82,976 | 15,025 | 76,104 | 3,871 | 0,667 | 122,885 | 16,826 | 72,993 | 5,818 | **0,019** | 0,666 |
| | 4M EFS 200 | 63,827 | 14,303 | 67,565 | 11,668 | 0,842 | 70,743 | 10,085 | 76,206 | 11,101 | 0,720 | 0,594 |
| | 4M EFS 400 | 63,568 | 14,458 | 70,230 | 9,817 | 0,708 | 70,603 | 9,899 | 77,240 | 10,367 | 0,650 | 0,63 |

### Table 6

| C-Off AT + G | | Control AT | | Control AT + G | | p | Rotenone AT | | Rotenone AT + G | | p | p Control vs Rotenone |
|---|---|---|---|---|---|---|---|---|---|---|---|---|
| | | Mean | SEM | Mean | SEM | | Mean | SEM | Mean | SEM | | |
| Duodenum | 2M EFS 200 | 79,173 | 14,143 | 90,052 | 7,056 | 0,507 | 73,393 | 11,249 | 103,054 | 14,259 | 0,133 | 0,433 |
| | 2M EFS 400 | 95,322 | 20,113 | 95,215 | 14,599 | 0,997 | 90,426 | 18,887 | 88,545 | 11,469 | 0,934 | 0,727 |
| | 4M EFS 200 | 81,353 | 9,268 | 79,459 | 9,085 | 0,886 | 65,884 | 6,538 | 74,802 | 6,934 | 0,363 | 0,686 |
| | 4M EFS 400 | 88,081 | 8,791 | 102,368 | 8,660 | 0,266 | 68,931 | 6,566 | 86,449 | 8,614 | 0,125 | 0,214 |
| Jejunum | 2M EFS 200 | 95,973 | 15,359 | 91,761 | 12,065 | 0,833 | 78,510 | 32,023 | 100,143 | 25,408 | 0,608 | 0,772 |
| | 2M EFS 400 | 98,213 | 15,010 | 97,778 | 8,622 | 0,981 | 75,032 | 14,189 | 82,999 | 9,473 | 0,651 | 0,275 |
| | 4M EFS 200 | 68,503 | 10,650 | 78,574 | 8,404 | 0,569 | 79,165 | 10,480 | 95,389 | 9,728 | 0,273 | 0,209 |
| | 4M EFS 400 | 74,035 | 7,839 | 90,882 | 16,642 | 0,274 | 82,448 | 10,150 | 98,868 | 8,893 | 0,241 | 0,612 |
| Ileum | 2M EFS 200 | 73,371 | 25,338 | 84,767 | 13,115 | 0,698 | 65,949 | 12,992 | 97,729 | 14,628 | 0,135 | 0,524 |
| | 2M EFS 400 | 80,838 | 20,061 | 91,997 | 12,890 | 0,650 | 76,370 | 10,713 | 105,610 | 12,336 | 0,104 | 0,463 |
| | 4M EFS 200 | 81,073 | 21,466 | 106,900 | 20,971 | 0,404 | 59,825 | 6,443 | 121,527 | 17,834 | **0,004** | 0,601 |
| | 4M EFS 400 | 76,077 | 14,726 | 107,241 | 15,902 | 0,172 | 63,085 | 6,077 | 116,604 | 24,860 | **0,004** | 0,673 |
| Colon | 2M EFS 200 | 99,826 | 36,553 | 190,669 | 32,531 | 0,093 | 27,022 | 11,878 | 234,174 | 39,160 | **<0,001** | 0,413 |
| | 2M EFS 400 | 94,823 | 47,659 | 564,570 | 365,080 | 0,231 | -16,350 | 27,606 | 1484,669 | 1165,451 | 0,227 | 0,468 |
| | 4M EFS 200 | 109,166 | 16,862 | 145,785 | 16,408 | 0,139 | 97,017 | 13,881 | 110,867 | 14,259 | 0,496 | 0,128 |
| | 4M EFS 400 | 104,363 | 11,605 | 132,025 | 14,554 | 0,157 | 97,498 | 16,565 | 120,712 | 17,657 | 0,352 | 0,628 |

### Table 7

| Serotonin max | Control | | Rotenone | |
|---|---|---|---|---|
| | Mean | SEM | Mean | SEM |
| Duodenum | 1,395 | 0,13 | 1,474 | 0,127 |
| Jejunum | 1,209 | 0,046 | 1,205 | 0,099 |
| Ileum | 1,009 | 0,031 | 0,924 | 0,035 |
| Colon | 0,877 | 0,055 | 0,851 | 0,042 |

| p | Jejunum | Ileum | Colon |
|---|---|---|---|
| Duodenum | 0,300 | **0,006** | **<0,001** |
| Jejunum | | 0,240 | **0,013** |
| Ileum | | | 0,594 |

### Table 8

| Serotonin min | Control | | Rotenone | |
|---|---|---|---|---|
| | Mean | SEM | Mean | SEM |
| Duodenum | 1,202 | 0,086 | 1,249 | 0,080 |
| Jejunum | 1,118 | 0,066 | 1,068 | 0,029 |
| Ileum | 0,933 | 0,033 | 0,914 | 0,055 |
| Colon | 1,030 | 0,085 | 0,972 | 0,026 |

| p | Jejunum | Ileum | Colon |
|---|---|---|---|
| Duodenum | 0,840 | **0,069** | 0,342 |
| Jejunum | | 0,284 | 0,807 |
| Ileum | | | 0,778 |

#### Table 9

| M3-Receptor | | Control | | Rotenone | | p |
|---|---|---|---|---|---|---|
| | | Mean | SEM | Mean | SEM | |
| Duodenum | 2M | 1 | 0,068 | 1,187 | 0,091 | **<0,01** |
| | 4M | 1 | 0,040 | 1,163 | 0,025 | **<0,01** |
| Jejunum | 2M | 1 | 0,033 | 1,224 | 0,046 | **<0,01** |
| | 4M | 1 | 0,033 | 1,151 | 0,068 | **<0,05** |
| Ileum | 2M | 1 | 0,057 | 1,229 | 0,059 | **<0,05** |
| | 4M | 1 | 0,062 | 1,223 | 0,035 | **<0,05** |
| Colon | 2M | 1 | 0,042 | 1,166 | 0,456 | **<0,05** |
| | 4M | 1 | 0,061 | 1,305 | 0,060 | **<0,01** |

#### Table 10

| ß2-subunit | | Control | | Rotenone | | p |
|---|---|---|---|---|---|---|
| | | Mean | SEM | Mean | SEM | |
| Duodenum | 2M | 1 | 0,105 | 1,260 | 0,038 | **<0,05** |
| | 4M | 1 | 0,012 | 1,425 | 0,104 | **<0,01** |
| Jejunum | 2M | 1 | 0,065 | 1,174 | 0,059 | >0,05 |
| | 4M | 1 | 0,033 | 1,809 | 0,240 | **<0,05** |
| Ileum | 2M | 1 | 0,040 | 1,181 | 0,091 | >0,05 |
| | 4M | 1 | 0,031 | 1,212 | 0,040 | **<0,01** |
| Colon | 2M | 1 | 0,012 | 1,287 | 0,067 | **<0,01** |
| | 4M | 1 | 0,084 | 1,570 | 0,122 | **<0,01** |

#### Table 11

| D2-Receptor | | Control | | Rotenone | | p |
|---|---|---|---|---|---|---|
| | | Mean | SEM | Mean | SEM | |
| Duodenum | 2M | 1 | 0,044 | 1,065 | 0,066 | >0,05 |
| | 4M | 1 | 0,190 | 1,653 | 0,189 | **<0,05** |
| Jejunum | 2M | 1 | 0,060 | 1,283 | 0,100 | **<0,05** |
| | 4M | 1 | 0,150 | 1,179 | 0,080 | >0,05 |
| Ileum | 2M | 1 | 0,005 | 1,151 | 0,078 | **<0,01** |
| | 4M | 1 | 0,047 | 1,153 | 0,060 | >0,05 |
| Colon | 2M | 1 | 0,064 | 1,514 | 0,016 | **<0,05** |
| | 4M | 1 | 0,046 | 1,205 | 0,029 | **<0,01** |

### Table 12

| PGP9.5 | | Control | | Rotenone | | p |
|---|---|---|---|---|---|---|
| | | Mean | SEM | Mean | SEM | |
| Duodenum | 2M | 1 | 0,17 | 0,60 | 0,09 | >0,05 |
| | 4M | 1 | 0,08 | 0,77 | 0,11 | >0,05 |
| Jejunum | 2M | 1 | 0,15 | 0,76 | 0,15 | >0,05 |
| | 4M | 1 | 0,03 | 1,01 | 0,28 | >0,05 |
| Ileum | 2M | 1 | 0,02 | 1,02 | 0,03 | >0,05 |
| Colon | 2M | 1 | 0,16 | 1,26 | 0,06 | >0,05 |
| | 4M | 1 | 0,31 | 1,01 | 0,11 | >0,05 |

### Table 13

| ChAT | | Control | | Rotenone | | p |
|---|---|---|---|---|---|---|
| | | Mean | SEM | Mean | SEM | |
| Duodenum | 2M | 1 | 0,11 | 0,84 | 0,16 | >0,05 |
| | 4M | 1 | 0,17 | 0,58 | 0,09 | >0,05 |
| Jejunum | 2M | 1 | 0,10 | 0,90 | 0,10 | >0,05 |
| | 4M | 1 | 0,26 | 0,58 | 0,07 | >0,05 |
| Colon | 2M | 1 | 0,15 | 0,57 | 0,06 | **<0,05** |
| | 4M | 1 | 0,26 | 2,00 | 0,04 | >0,05 |
| All regions | 2M | 1 | 0,06 | 0,77 | 0,06 | **<0,05** |
| | 4M | 1 | 0,08 | 0,75 | 0,06 | **<0,05** |

### Table 14

| TH | | Control | | Rotenone | | p |
|---|---|---|---|---|---|---|
| | | Mean | SEM | Mean | SEM | |
| Duodenum | 2M | 1 | 0,07 | 1,71 | 0,15 | **<0,01** |
| | 4M | 1 | 0,17 | 0,78 | 0,19 | >0,05 |
| Jejunum | 2M | 1 | 0,05 | 0,66 | 0,13 | **<0,05** |
| | 4M | 1 | 0,23 | 0,98 | 0,24 | >0,05 |
| Ileum | 2M | 1 | 0,09 | 1,07 | 0,23 | >0,05 |
| Colon | 2M | 1 | 0,10 | 1,48 | 0,14 | **<0,05** |
| | 4M | 1 | 0,24 | 1,29 | 0,16 | >0,05 |
| All regions | 2M | 1 | 0,06 | 0,78 | 0,06 | >0,05 |
| | 4M | 1 | 0,08 | 0,75 | 0,06 | >0,05 |

**Table 1 Statistical results comparing the reaction to carbachol between vehicle and rotenone-exposed mice.** A 2-way ANOVA was used to compare the effect of carbachol on the amplitude, the maximal exerted contraction force and the muscle tone between vehicle and rotenone-exposed mice. The level of significance is shown as the p value.

**Table 2 Statistical results comparing the reaction to dopamine between vehicle and rotenone-exposed mice.** A 2-way ANOVA was used to compare the effect of dopamine on the amplitude, the maximal exerted contraction force and the muscle tone between vehicle and rotenone-exposed mice. The level of significance is shown as the p value.

**Table 3** shows the mean values (mean), the standard error of the mean (SEM) and the significance (p) of the comparison (unpaired Student´s t-test) between the relaxation observed in R1 induced by the EFS (200 and 400 m µs) in the absence of any blocker (consider as 100%) and the relaxation obtained in the presence of atropine (AT) normalized to the value without substance and shown as percentage. The far-right column shows the significance when comparing the results obtained in the presence of AT between control and rotenone exposed mice.

**Table 4** shows the mean values (mean), the standard error of the mean (SEM) and the significance (p) of the comparison (unpaired Student´s t-test) between the contraction observed in C-Off induced by the EFS (200 and 400 µs) in the absence of any blocker (consider as 100%) and the contraction obtained in the presence of atropine (AT). The far-right column shows the significance when comparing the results obtained in the presence of AT between control and rotenone exposed mice.

**Table 5** shows the mean values (mean), the standard error of the mean (SEM) and the significance (p) of the comparison (unpaired Student´s t-test) between the relaxation (R1) obtained with the EFS (200 and 400 µs) in the presence of atropine (AT) and in the presence of AT and guanethidine (G). The absence of any blocker is considered as 100% and all other values were normalized to that value. The far-right column shows the significance when comparing the results obtained in the presence of AT and G between control and rotenone exposed mice.

**Table 6** shows the mean values (mean), the standard error of the mean (SEM) and the significance (p) of the comparison (unpaired Student´s t-test) between the contraction (C-Off) obtained with the EFS (200 and 400 µs) in the presence of atropine (AT) and in the presence of AT and guanethidine (G). The absence of any blocker is considered as 100% and all other values were normalized to that value.

The far-right column shows the significance when comparing the results obtained in the presence of AT and G between control and rotenone exposed mice.

**Table 7** shows the mean values (mean) and the standard error of the mean (SEM) for maximal tension exerted by serotonin (S) when added in the presence of AT and G. The *P*-value table shows the significance of the difference in the effect that S had between regions of the intestine.

**Table 8** shows the mean values (mean) and the standard error of the mean (SEM) for the muscle tone (Min.) by serotonin (S) when added in the presence of AT and G. The *P*-value table shows the significance of the difference in the effect that S had between regions of the intestine.

**Table 9** shows the mean values (mean), the standard error of the mean (SEM) and the significance (p) of the difference between control and rotenone exposed mice in the expression of the muscarinic cholinergic Receptor (M3) normalized to the total amount of neurons (PGP9.5) and to the mean of these values in the control group.

**Table 10** shows the mean values (mean), the standard error of the mean (SEM) and the significance (p) of the difference between control and rotenone exposed mice in the expression of the adrenergic Receptor (ß2-subunit) normalized to the total amount of neurons (PGP9.5) and to the mean of these values in the control group.

**Table 11** shows the mean values (mean), the standard error of the mean (SEM) and the significance (p) of the difference between control and rotenone exposed mice in the expression of the dopaminergic Receptor (D2) normalized to the total amount of neurons (PGP9.5) and to the mean of these values in the control group.

**Table 12** shows the mean values (mean), the standard error of the mean (SEM) and the significance (p) of the difference between control and rotenone exposed mice in the expression of PGP 9.5 normalized to the mean of the control group.

**Table 13** shows the mean values (mean), the standard error of the mean (SEM) and the significance (p) of the difference between control and rotenone exposed mice in the expression of choline acetyltransferase (ChAT) normalized to the total amount of neurons (PGP9.5) and to the mean of these values in the control group.

**Table 14** shows the mean values (mean), the standard error of the mean (SEM) and the significance (p) of the difference between control and rotenone exposed mice in the expression of tyrosine hydroxylase (TH) normalized to the total amount of neurons (PGP9.5) and to the mean of these values in the control group.
