## Supplementary material for "Physiological and pathological changes in the enteric nervous system of rotenone-exposed mice as early biomarkers for Parkinson’s disease": Quantification code MatLab

function paco5

% letzte ─nderung: K.B. 10.01.2017

```
close all
clear


global setup ftsize b a b2 a2 erg


% Think pad : FontSize',12,
% GNR6PN:     FontSize',10,   screen size: 1 1 1024 768

ftsize = 12;

scrsz = get(0,'ScreenSize');
f = figure('Position',[1,20,scrsz(3), scrsz(4)*0.96],'Toolbar','none',...
                'MenuBar','none');
```

setup.ErgFilename = 'PacoErgebnis.txt'; % Hier den Dateinamen Σndern auf dem die Ergebnisse gespeichert weren (.txt) !

setup.ErgFilepath = 'D:\A-Data\Paco\Auswertung'; % Hier den Ordner Σndern, in dem die Ergebnis-Datei steht

```
setup.DataFilename = ' ';
setup.DataFilepath = 'D:\A-Data\Paco';                                  % Hier den Ordner eintragen, in dem nach Daten gesucht werden soll


uicontrol('Style','pushbutton','String','Input-Datei',...
          'Units','normalized',...
          'Position',[0.05, 0.03, 0.06, 0.03],...
          'Callback',@ReadData_Callback);
```

uicontrol('Style','pushbutton','String','auswΣhlen',... % Zeitbereich auswΣhlen

'Units','normalized',...

'Position',[0.12, 0.03, 0.06, 0.03],...

'Callback',@select_Callback);

uicontrol('Style','pushbutton','String','alles zeigen',... % Zeitbereich auswΣhlen

'Units','normalized',...

'Position',[0.19, 0.03, 0.06, 0.03],...

'Callback',@showall_Callback);

uicontrol('Style','pushbutton','String','<',... % Zeitbereich auswΣhlen

'Units','normalized',...

'Position',[0.26, 0.03, 0.03, 0.03],...

'Callback',@back_Callback);

uicontrol('Style','pushbutton','String','>',... % Zeitbereich auswΣhlen

```
          'Units','normalized',...
          'Position',[0.31, 0.03, 0.03, 0.03],...
          'Callback',@forward_Callback);

uicontrol('Style','pushbutton','String',' >>',...                      % next com
          'Units','normalized',...
          'Position',[0.36, 0.03, 0.03, 0.03],...
          'Callback',@next_Callback);

uicontrol('Style','pushbutton','String',' messen',...                      % next com
          'Units','normalized',...
          'Position',[0.41, 0.03, 0.06, 0.03],...
          'Callback',@messen_Callback);
```

uicontrol('Style','text','String','Pre-time (min):',... % pretime auswΣhlen

'Units','normalized','BackgroundColor',[0.8 0.8 0.8],...

'Position',[0.75, 0.024, 0.07, 0.03]);

uicontrol('Style','edit','String','1',... % pretime auswΣhlen

'Units','normalized',...

'Position',[0.82, 0.03, 0.03, 0.03],...

'Callback',@pretime_Callback);

uicontrol('Style','text','String','Post-time (min):',... % pretime auswΣhlen

'Units','normalized','BackgroundColor',[0.8 0.8 0.8],...

'Position',[0.86, 0.024, 0.07, 0.03]);

uicontrol('Style','edit','String','1',... % pretime auswΣhlen

```
          'Units','normalized',...
          'Position',[0.93, 0.03, 0.03, 0.03],...
          'Callback',@posttime_Callback);

uicontrol('Style','pushbutton','String','Markierung versetzen',...                      % next com
          'Units','normalized',...
          'Position',[0.55, 0.03, 0.08, 0.03],...
          'Callback',@Markierung_versetzen_Callback);

uicontrol('Style','pushbutton','String','Grafik von diesem Abschnitt',...                      % next com
          'Units','normalized',...
          'Position',[0.65, 0.03, 0.1, 0.03],...
          'Callback',@Grafik_Abschnitt_Callback);

f2 = figure('Position',[20,90,scrsz(3)*0.95, scrsz(4)*0.8],'Toolbar','none',...
                'MenuBar','none','vis','off');

uicontrol('Style','pushbutton','String',' weiter ohne speichern',...                      % next com
          'Units','normalized',...
          'Position',[0.05, 0.03, 0.2, 0.03],...
          'Callback',@weiter_ohne_speichern_Callback);
uicontrol('Style','pushbutton','String',' Anstieg messen',...                      % next com
          'Units','normalized',...
          'Position',[0.35, 0.03, 0.2, 0.03],...
          'Callback',@anstieg_messen_Callback);


uicontrol('Style','pushbutton','String',' weiter mit speichern',...                      % next com
          'Units','normalized',...
          'Position',[0.7, 0.03, 0.2, 0.03],...
          'Callback',@weiter_mit_speichern_Callback);
```

message = uicontrol('Style','text','String','Beginn des Anstiegs markieren',... % pretime auswΣhlen

```
          'Units','normalized','FontSize',12,'vis','off',...
          'Position',[0.35, 0.2, 0.3, 0.04]);

%  f3 = figure('Position',[40,90,scrsz(3)*0.9, scrsz(4)*0.5],'vis','off');

extra_figures = cell(1,5);

extra_figures{1} = figure('Position',[20,90,scrsz(3)*0.2, scrsz(4)*0.5],'vis','off');
extra_figures{2} = figure('Position',[scrsz(3)*0.23,90,scrsz(3)*0.2, scrsz(4)*0.5],'vis','off');
extra_figures{3} = figure('Position',[scrsz(3)*0.45,90,scrsz(3)*0.2, scrsz(4)*0.5],'vis','off');
extra_figures{4} = figure('Position',[scrsz(3)*0.67,90,scrsz(3)*0.2, scrsz(4)*0.5],'vis','off');
extra_figures{5} = figure('Position',[scrsz(3)*0.89,90,scrsz(3)*0.05, scrsz(4)*0.5],'vis','off');

uicontrol('Style','pushbutton','String',' weiter',...                      % next com
          'Units','normalized',...
          'Position',[0.05, 0.4, 0.9, 0.3],...
          'Callback',@weiter);
```

% uicontrol('Style','pushbutton','String','3',... % posttime auswΣhlen

```
%           'Units','normalized',...
%           'Position',[0.31, 0.03, 0.03, 0.03],...
%           'Callback',@posttime_Callback);


      % 'FontSize',ftsize,'FontName','Times','FontWeight','bold',...

% d =
%
%                  data: [1x34006016 double]
%             datastart: [8x2 double]
%               dataend: [8x2 double]
%                titles: [8x10 char]
%              rangemin: [8x2 double]
%              rangemax: [8x2 double]
%              unittext: 'mN'
%           unittextmap: [8x2 double]
%            blocktimes: [7.3452e+05 7.3452e+05]
%              tickrate: [400 400]
%            samplerate: [8x2 double]
%     firstsampleoffset: [8x2 double]
%               comtext: [17x18 char]
%                   com: [31x5 double]   Spalte 3: Zeit, Spalte 5 Typ (bezieht sich auf comtext)


function ReadData_Callback(~, ~)
    [setup.DataFilename,setup.DataFilepath] = uigetfile([setup.DataFilepath,'\*.mat'], 'Select *.mat - file');
    d=load(fullfile(setup.DataFilepath,setup.DataFilename));


    % Daten werden verkleinert, nur jeder 8. Wert wird genommen, da sonst das
    % Programm zu langsam ist
    % Zeitachse in Minuten, beginnt fⁿr jeden neuen versuch wieder bei Null
```

[setup.Nkanal, Nversuch] = size(d.datastart); % Anzahl von KanΣlen und Versuchen

d.data = d.data(1:setup.Nkanal:end);

d.datastart = (d.datastart+7) / setup.Nkanal;

d.dataend = ((d.dataend+setup.Nkanal) / setup.Nkanal) -1;

d.samplerate = mean(d.samplerate) / setup.Nkanal;

d.com(:,3) = round(d.com(:,3)/setup.Nkanal); % Indices an denen ein Kommentar steht Hier als zeitwert in Minuten

data_length = mean(diff(d.datastart)); % LΣnge des Datenvektors eines Kanals in jedem Experiment

```
    % if Nversuch == 2, vers = 2; else vers = 1; end                           % Hier wird davon ausgegangen, dass immer der letzte Versuch wichtig ist
    vers = Nversuch;

    setup.samplerate = d.samplerate(vers);
    setup.time = (0:data_length(vers)-1) ./ setup.samplerate ./ 60;           % time in Min.   Samplerate in samples pro s !!!!
    an=d.datastart(1,vers);                                             % an und en sind immer Indices, nie Zeitwerte
    en = d.dataend(setup.Nkanal,vers);
    setup.data = reshape(d.data(an:en)',data_length(vers),setup.Nkanal);
    setup.com = d.com(d.com(:,2) == vers,:);
    setup.comtext = d.comtext;
    setup.pre_post = [1 1 0 0];                                         % Zeit in Min (1+2) und in datenpunkten (3+4)
    setup.pre_post(3:4) = round(setup.pre_post(1:2) .* setup.samplerate .* 60);
    setup.an = 1;
    setup.en = data_length(vers);
    setup.next_comment = 0;
    setup.titles = d.titles;
    setup.chan = 0;                                                         % aktueller Kanal, der gemessen wird


    erg = zeros(10,setup.Nkanal);                  % Ergebnisse, fⁿr jees ereignis ein Ergebnisblock, wird als textfile gespeichert

    upper_freq_limit=0.2;
    [b,a]=butter(3,upper_freq_limit*2/d.samplerate(vers));

    upper_freq_limit=2 ;
    [b2 a2]=butter(3,upper_freq_limit*2/d.samplerate(vers));


    axes('position', [0 0 1 1],'visible','off')
    txt = sprintf('File: %s   Gesamtdauer: %5.2f Min. ', setup.DataFilename, setup.time(end));
    text(0.5 ,0.99,txt,'units','normalized','horizontalAlignment','center','VerticalAlignment','top','Fontsize',12)


    fprintf(' \n\n')
    [r,~] = size(setup.comtext);
    for n=1:r
        fprintf('%d) %s \n',n,setup.comtext(n,:))
    end
    fprintf(' \n\n')

    setup.com

    plot_all
end

    function select_Callback(~,~)
        [x,~] = ginput(2);
        [~,setup.an] = min(abs(setup.time-x(1)));
        [~,setup.en] = min(abs(setup.time-x(2)));
        plot_all
    end
function showall_Callback(~,~)
        setup.an = 1;
        setup.en = length(setup.time);
        setup.next_comment = 0;
        plot_all
end
function back_Callback(~,~)
        dummy = setup.en - setup.an;
        setup.an = setup.an - dummy;
        if setup.an < 1
            setup.an = 1;
        end
        setup.en = setup.an + dummy;

        plot_all
end
function forward_Callback(~,~)
        dummy = setup.en - setup.an;
        setup.an = setup.an + dummy;
        setup.en = setup.an + dummy;

        if setup.en > length(setup.time)
            setup.en = length(setup.time);
            setup.an = setup.en - dummy;
         end

        plot_all
end

function pretime_Callback(src,~)
    setup.pre_post(1) = str2double(get(src,'String'));
    setup.pre_post(3) = round(setup.pre_post(1) .* setup.samplerate .* 60);
    setup.an=setup.com(setup.next_comment,3) - setup.pre_post(3);                     % index begin next window
    plot_all
end
function posttime_Callback(src,~)
    setup.pre_post(2) = str2double(get(src,'String'));
    setup.pre_post(4) = round(setup.pre_post(2) .* setup.samplerate .* 60);
    setup.en=setup.com(setup.next_comment,3) + setup.pre_post(4);                     % index begin next window
    plot_all
end
function next_Callback(~,~)                                                            % go to next comment/event
    if setup.next_comment < length(setup.com)
       setup.next_comment = setup.next_comment + 1;
       setup.an=setup.com(setup.next_comment,3) - setup.pre_post(3);                     % index begin next window
       setup.en=setup.com(setup.next_comment,3) + setup.pre_post(4);                     % index begin next window
       fprintf('setup.next_comment = %d; setup.an = %d\n', setup.next_comment, setup.an)
    else
       setup.an = 1;
       setup.en = length(setup.data);
       setup.next_comment = 0;                                                     % reset
    end
    plot_all
    t=setup.com(setup.next_comment,5);            % typ comment
    axes('position', [0 0 1 1],'visible','off');
    txt = sprintf('Ereignis Nr %d/%d Typ %d (%s)', setup.next_comment, length(setup.com),t,deblank(setup.comtext(t,:)));
    text(0.5 ,0.96,txt,'units','normalized','horizontalAlignment','center','VerticalAlignment','top','Fontsize',10)
end


% function Markierung_versetzen_Callback(~,~)
%      marker_ind = find((setup.com(:,3) > setup.an) & (setup.com(:,3) < setup.en));
%      if length(marker_ind) == 1
%          [x,~] = ginput(1);
%          [~,ind] = min(abs(setup.time-x));
% %          fprintf('Neuer Marker bei %0.6f \n',setup.time(ind))
%            setup.com(marker_ind,3) = ind;
%            plot_all
%      end
% end

function Markierung_versetzen_Callback(~,~)
       [x,~] = ginput(1);                                     % der erste click geht auf die Markierung, die ersetzt werden soll
       [~,ind] = min(abs(setup.time-x));                      % ind ist der Index
       [~,marker_ind] = min(abs(setup.com(:,3)-ind));                % ind ist jetzt der Index des Markers
       [x,~] = ginput(1);                                     % jetzt auf die neue Position klicken
       [~,ind] = min(abs(setup.time-x));
       setup.com(marker_ind,3) = ind;                         % jetzt hat der Marker eine neue Position
       plot_all
end


function Grafik_Abschnitt_Callback(~,~)
    an = setup.an;
    en = setup.en;
    ti = (0:(setup.en - setup.an)) ./ setup.samplerate ./ 60;
    t=setup.com(setup.next_comment,5);
        for n=1:4
           set(extra_figures{n},'vis','on')
           subplot(2,1,1)
           plot(ti,setup.data(an:en,n),'k','LineWidth',1.25)
           axis([ti(1) ti(end) min(setup.data(an:en,n)) min(setup.data(an:en,n)) + 1.2 * (max(setup.data(an:en,n)) - min(setup.data(an:en,n)))]);
           ylabel('[ mN ]')
           set(gca,'box','off','TickDir','out','LineWidth',1.25)
           title(setup.titles(n,:))
           t1 = deblank(setup.comtext(t,:));
           ind = strfind(t1,'-');
           if ~isempty(ind)
               t1 = [t1(1:ind-1),'^{', t1(ind:ind+2),'}',t1(ind+3:end)];
           end
           txt = sprintf('%s, Control', t1);
           text(0.5,0.9,txt,'HorizontalAlignment','center','units','normalized','FontName','Arial')

           subplot(2,1,2)
           plot(ti,setup.data(an:en,n+4),'k','LineWidth',1.25)
           axis([ti(1) ti(end) min(setup.data(an:en,n+4)) min(setup.data(an:en,n+4)) + 1.2 * (max(setup.data(an:en,n+4)) - min(setup.data(an:en,n+4)))]);
           ylabel('[ mN ]')
           set(gca,'box','off','TickDir','out','LineWidth',1.25)
           xlabel('Time [ Min ]')
           txt = sprintf('%s, 5mg/kg rotenone', t1);
           text(0.5,0.9,txt,'HorizontalAlignment','center','units','normalized','FontName','Arial')
        end
        set(extra_figures{5},'vis','on')
end


function messen_Callback(~,~)                                                   % missst einen Kanal  aus
    setup.chan = setup.chan + 1;
    if setup.chan > setup.Nkanal
        setup.chan = 0;
        erg(:,:) = 0;                                                           % Ergebnisse resetten
        set(f2,'vis','off')
    else                                                                        % jetzt ausmesssen
        an = setup.an;
        en = setup.en;
        marker_ind = setup.com(setup.next_comment,3);                           % Index des markers, um den gemessen wird
        set(f2,'vis','on')
        plot(setup.time(an:en),setup.data(an:en,setup.chan))                         % Daten plotten
        set(gca,'position', [0.06 0.3 0.9 0.6])
        axis tight

        ax=axis;                                             % vertikale Linie bei Ereignis
        y = ax(3) + (ax(4)-ax(3)) * 0.85;
        line([setup.time(marker_ind) setup.time(marker_ind)],[ax(3) ax(4)])
        text(setup.time(marker_ind), y, num2str(setup.com(setup.next_comment,5)))

        title([num2str(setup.chan),'.) ', setup.titles(setup.chan,:)])
        xlabel('Zeit [ Min ]')

        temp_data = filtfilt(b,a,setup.data(an:en,setup.chan));                  % mittelwerte rot plotten
        line(setup.time(an:en),temp_data,'color',[1,0,0],'linewidth',1.5)

        temp_data = filtfilt(b2,a2,setup.data(an:en,setup.chan));                % Minima und Maxima finden
        temp_data=diff(temp_data);
        ind1 = find(temp_data>0);
        ind2 = find(diff(ind1) > 1);
        maxima = ind1(ind2)+1+an-1;                                              % indices beziehen sich auf setup.data
        minima = ind1(ind2+1)+an-1;                                              % indices beziehen sich auf setup.data

        ind1 = find(minima <= marker_ind);                        % indices maxima und minima prae
        ind2 = find(minima > marker_ind);                         % indices maxima und minima post

        hold on
        plot(setup.time(maxima(ind1)),setup.data(maxima(ind1),setup.chan),'ro')
        plot(setup.time(minima(ind1)),setup.data(minima(ind1),setup.chan),'ro')

        plot(setup.time(maxima(ind2)),setup.data(maxima(ind2),setup.chan),'go')
        plot(setup.time(minima(ind2)),setup.data(minima(ind2),setup.chan),'go')
        hold off

        erg(setup.chan,1) = mean(setup.data(maxima(ind1),setup.chan) - setup.data(minima(ind1),setup.chan));              % Erg 1 : mean Amplitude prae
        erg(setup.chan,2) = mean(setup.data(maxima(ind2),setup.chan) - setup.data(minima(ind2),setup.chan));              % Erg 2 : mean Amplitude post
        erg(setup.chan,3) = max(setup.data(maxima(ind1),setup.chan));                                     % Erg 3 : Maximum prae
        erg(setup.chan,4) = min(setup.data(minima(ind1),setup.chan));                                     % Erg 4 : Minimum prae
        erg(setup.chan,5) = max(setup.data(maxima(ind2),setup.chan));                                     % Erg 5 : Maximum post
        erg(setup.chan,6) = min(setup.data(minima(ind2),setup.chan));                                     % Erg 6 : Minimum post
        erg(setup.chan,7) = length(ind1) / (setup.time(marker_ind) - setup.time(an)) ;         % Erg 7 : Frequenz prae in Kontraktionen / Minute
        erg(setup.chan,8) = length(ind2) / (setup.time(en) - setup.time(marker_ind)) ;         % Erg 8 : Frequenz post in Kontraktionen / Minute
        fprintf('Messwerte: %10.3f /n',erg(setup.chan,1:8))
    end

end


function weiter_ohne_speichern_Callback(~,~)
    messen_Callback;
end

function weiter(~,~)
    for n=1:5
        set(extra_figures{n},'vis','off');
    end
%     plot_all;
end

function weiter_mit_speichern_Callback(~,~)
    messen_Callback;
end

function anstieg_messen_Callback(~,~)                                        % Beginn und Ende des steilen Anstiegs nach Ereignis soll markiert werden
    an = setup.an;
    en = setup.en;
    temp_data = filtfilt(b,a,setup.data(an:en,setup.chan));                  % mittelwerte rot plotten

    set(message,'vis','on','string','Beginn des Anstiegs markieren');
    [x,~] = ginput(1);
    [~,ind1] = min(abs(setup.time-x));  ind1=ind1-an+1;                   % ind1 (Beginn des Anstiegs) bezieht sich jetzt auf temp_data

    set(message,'vis','on','string','Maximum des Anstiegs markieren');
    [x,~] = ginput(1);
    [~,ind2] = min(abs(setup.time-x));  ind2=ind2-an+1;

    set(message,'vis','on','string','Ende und Plateau des Abfalls markieren');
    [x,offset] = ginput(1);
    [~,ind3] = min(abs(setup.time-x));  ind3=ind3-an+1;
    set(message,'vis','off');

    x=setup.time((ind1:ind2)+an-1); x = x(:);
    y = temp_data(ind1:ind2); y=y(:);
    const = ones(ind2-ind1+1,1);
    Coeffs = [x,const] \ y;       %Fit line to Anstieg
    erg(setup.chan,9) = Coeffs(1);                                           % Anstigssteilheit
    % erg(setup.chan,10) = Coeffs(2);
    y = x .* Coeffs(1) + Coeffs(2);
    line(x,y,'color',[0 0 0],'linewidth',1.5)
    erg(setup.chan,11) = temp_data(ind2) - temp_data(ind1);                  % Anstiegsamplitude

                                                                               % Hinten muss eine Nullinie sein zum fitten der e-funktion
    y = temp_data(ind2:ind3)-offset; y=y(:);                                 % abfallende Flanke mit exponential fit
    x = setup.time(1:length(y));x=x(:);                                     % Zeitachse
    ff = fit(x,y,'exp1');
%      plot(ff,x,y)
    fit_data = ff.a * exp(ff.b*x);
   line(setup.time((ind2:ind3)+an-1),fit_data + offset,'color',[0 0 0],'linewidth',1.5)
%     [~,~] = ginput(1);


end


function plot_all
    an = setup.an;
    en = setup.en;
    marker_ind = find((setup.com(:,3) > an) & (setup.com(:,3) < en));           % index an denen ein event ist
    fprintf('an = %d; en = %d\n',an,en)

    for n=1:setup.Nkanal
        subplot(setup.Nkanal/2,2,n)
        plot(setup.time(an:en),setup.data(an:en,n))
        axis tight
        ax=axis;
        y = ax(3) + (ax(4)-ax(3)) * 0.85;
        if ~isempty(marker_ind)
            for n2=marker_ind'
                line([setup.time(setup.com(n2,3)) setup.time(setup.com(n2,3))],[ax(3) ax(4)])
                text(setup.time(setup.com(n2,3)), y, num2str(setup.com(n2,5)))
            end
        end
        title([num2str(n),'.) ', setup.titles(n,:)])
        if n > (setup.Nkanal-2)
            xlabel('Zeit [ Min ]')
        end
    end


    axes('position', [0 0 1 1],'visible','off');
    txt = sprintf('File: %s   Gesamtdauer: %5.2f Min.', setup.DataFilename, setup.time(end));
    text(0.5 ,0.99,txt,'units','normalized','horizontalAlignment','center','VerticalAlignment','top','Fontsize',12)

end

end


% Atomzerfall:
%
% N=N0 * exp(-lambda*t)
% hier
% y=b + N0 * exp(-lambda*(t-t0))

% Matlab:
% y = a * exp(bx)
```
